# Aberrant paracrine signalling for bone remodelling underlies the mutant histone-driven giant cell tumour of bone

**DOI:** 10.1101/2021.12.27.474195

**Authors:** Lucia Cottone, Lorena Ligammari, Helen J. Knowles, Hang-Mao Lee, Stephen Henderson, Sara Bianco, Christopher Davies, Sandra Strauss, Fernanda Amary, Ana Paula Leite, Roberto Tirabosco, Kristian Haendler, Joachim L. Schultze, Javier Herrero, Paul O’Donnell, Agamemnon E. Grigoriadis, Paolo Salomoni, Adrienne M. Flanagan

## Abstract

Oncohistones represent compelling evidence for a causative role of epigenetic perturbations in cancer. Giant cell tumours of bone (GCTs) are characterised by a mutated histone H3.3 as the sole genetic driver present in bone-forming osteoprogenitor cells but absent from abnormally large bone-resorbing osteoclasts which represent the hallmark of these neoplasms. While these striking features imply a pathogenic interaction between mesenchymal and myelomonocytic lineages during GCT development, the underlying mechanisms remain unknown.

We show that the changes in the transcriptome and epigenome in the mesenchymal cells caused by the H3.3-G34W mutation contribute to increase osteoclast recruitment in part via reduced expression of the TGFβ-like soluble factor, SCUBE3. In turn, osteoclasts secrete unregulated amounts of SEMA4D enhancing proliferation of mutated osteoprogenitors and arresting their maturation. These findings provide a mechanism by which GCTs undergo differentiation upon denosumab treatment, a drug that depletes osteoclasts. In contrast, gain of *hTERT* activity, commonly found in malignant GCT, makes neoplastic cells insensitive to osteoclasts, predicting the unresponsiveness to denosumab.

We provide a mechanism for GCT initiation and its response to current treatment, the basis of which is dysfunctional cross-talk between bone-forming and bone-resorbing cells, emphasising the importance of tumor/microenvironment bidirectional interactions in tumorigenesis.

## Introduction

Giant cell tumour of bone (GCT) is a locally aggressive primary neoplasm of bone (1). At the genetic level it is characterised by the presence of a near universal H3.3 Histone A (*H3-3A*) G34W missense mutation (H3.3^G34W^) which represents the sole genetic driver (2). Transformation of conventional to malignant GCT requires acquisition of at least one additional driver alteration, commonly in Telomerase Reverse Transcriptase (*hTER*T), reflected in histological characteristics of high grade sarcoma (3).

GCT is composed of H3.3^G34W^-mutated osteoprogenitors/mesenchymal stromal bone-forming cells and a pronounced tumour microenvironment (TME) dominated by unusually large unmutated bone-resorbing osteoclasts containing up to 100 nuclei (2). The mechanisms that lead to such a conspicuous osteoclast population have not yet been elucidated, although there are reports implicating Receptor Activator of Nuclear Factor kappa-B Ligand (RANKL) (4,5) on which osteoclasts, cells of the myeloid lineage, depend for their formation (6).

Denosumab, a humanised antibody to RANKL which blocks osteoclast formation, is widely employed to reduce excessive bone resorption in osteoporosis and localised osteolysis associated with metastatic cancer and is also used to control growth of GCT by targeting the TME (4).

Histone H3.3 is a replication-independent variant histone which facilitates transcription at euchromatic regions (7). In addition to GCT, mutations in the histone H3.3 have been identified in gliomas (H3.3K27M or H3.3G34R/V) (8) and in chondroblastomas (H3.3K36M) (2). Recent evidence suggests that H3.3^G34W^ promotes PRC2/H3K27me3 silencing of H3K36me3-depleted nucleosomes, H3.3 redistribution (9) and changes in the DNA methylation profile in osteoprogenitors (5). This epigenetic remodelling is reported to alter mesenchymal lineage commitment, including stalling of the osteogenic differentiation process (5,9,10). However, these studies do not fully explain how mutated osteoblasts influence osteoclastogenesis, a key pathogenic component of oncohistone-driven bone neoplasms.

## Results

### H3.3^G34W^ in osteoprogenitors regulates bone formation without affecting proliferation

Clinical evidence shows that treatment of GCT with denosumab not only results in depletion of osteoclasts but also causes a reduction in proliferation and an increase in differentiation/bone formation of the mutant osteoprogenitors (11) (**Figure 1A-B**). This led us to hypothesise that the GCT is driven by the H3.3^G34W^-mutant osteoprogenitors in a non-cell autonomous manner by recruiting large osteoclasts in the TME and co-opting them to secrete factors which provide the growth advantage to the tumour cells (**Figure 1C**).

**Figure 1.**
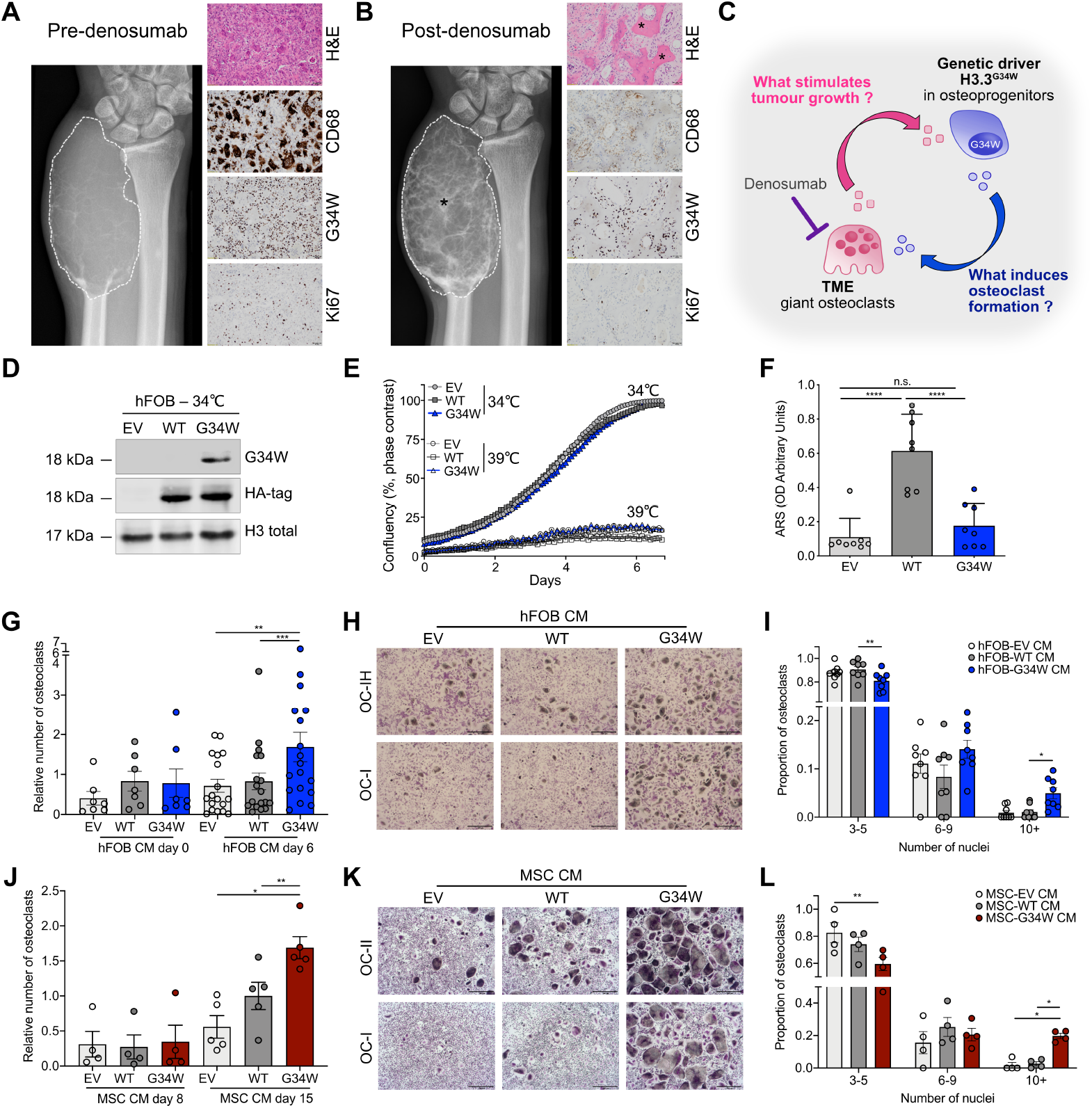
Mutant H3.3^G34W^ in osteoprogenitors regulates bone formation and stimulates osteoclast recruitment. A-B. GCT of the distal ulna in a 35-year-old male. (A) Pre-denosumab: anteroposterior radiograph of the distal ulna shows the tumour (dotted outline) without mineralisation. H&E-stained section: features of benign GCT with CD68-positive-osteoclasts interspersed with proliferating H3.3^G34W^-mononuclear stromal cells (G34W; Ki67). (B) Post-denosumab: after one month’s treatment showing similar tumour size (dotted outline) but prominent mineralisation (asterisk). H&E-section: bone formation (asterisk) and absence of CD68-positive-osteoclasts but persistent H3.3^G34W^-neoplastic cells (G34W) with reduced proliferative index (Ki67) compared to (A). **C**. Schema of proposed interactions between stromal/osteoprogenitors and TME. H3.3^G34W^-osteoprogenitor-derived factor(s) result in an environment permissive for unregulated formation of abnormally large osteoclasts, which secrete factor(s) stimulating tumour growth. Denosumab treatment inhibits osteoclasts removing the growth stimulus for stromal/osteoprogenitors. **D**. Western blot of H3.3^G34W^ expression in transfected hFOBs. **E**. Proliferation of undifferentiated hFOBs grown at 34ºC and differentiated at 39ºC for 6 days in mineralisation medium assessed by Incucyte; three replicates, two experiments. **F**. Quantification of ARS mineralisation of hFOBs on day 6. 8 replicates, three experiments. **G** and **J**. Number of osteoclasts generated in the presence of conditioned medium (CM) from hFOB (G) undifferentiated (day 0, 34 ºC) and differentiated at 39ºC for 6 days and (J) from iPSC-derived-MSCs differentiated to osteoblasts for 8 and 15 days; 7-18 (G) and 4-5 (J) preparations. **H** and **K**. Representative tartrate-resistant acid phosphatase (TRAP) staining of two osteoclast cultures in the presence of CM from (H) hFOBs differentiated for 6 days or (K) iPSC-derived-MSCs differentiated to osteoblasts for 15 days; 8 preparations. 4X magnification. **I** and **L**. Quantification of number of nuclei per osteoclast in (H) and (K); 4 preparations. Data are mean+/-SD (F), +/-SEM (G,J,I,L). F:1-way ANOVA. G,J: 1-way repeated measures (RM)ANOVA for each time point. I,L: 2-way RM ANOVA.

To elucidate the role of the H3.3^G34W^ mutation in the pathogenesis of GCT, we stably expressed H3.3^G34W^, H3.3^WT^ and empty vector (EV) *in vitro* in an immortalised human fetal osteoprogenitor cell line (hFOB) (**Figure 1D** and **Supplementary Figure 1A-B**).

Overexpression of H3.3^G34W^ did not alter cell proliferation, migration or survival compared to H3.3^WT^ or EV, even after long-term passaging (**Figure 1E** and **Supplementary Figure 1C-F**). Differentiation assays showed that while H3.3^WT^ promoted bone formation, H3.3^G34W^ impaired this function (**Figure 1F** and **Supplementary Figure 1G-H**). Overexpression of H3.3^G34W^ also had no effect on proliferation or survival of mesenchymal stem cells (MSCs) generated from induced Pluripotent Stem Cells (iPSCs) (**Supplementary Figure 2A-M**). Together, these results suggest that while H3.3^G34W^ does not directly control the growth of the stromal/osteoprogenitor cells, it regulates their differentiation.

**Figure 2.**
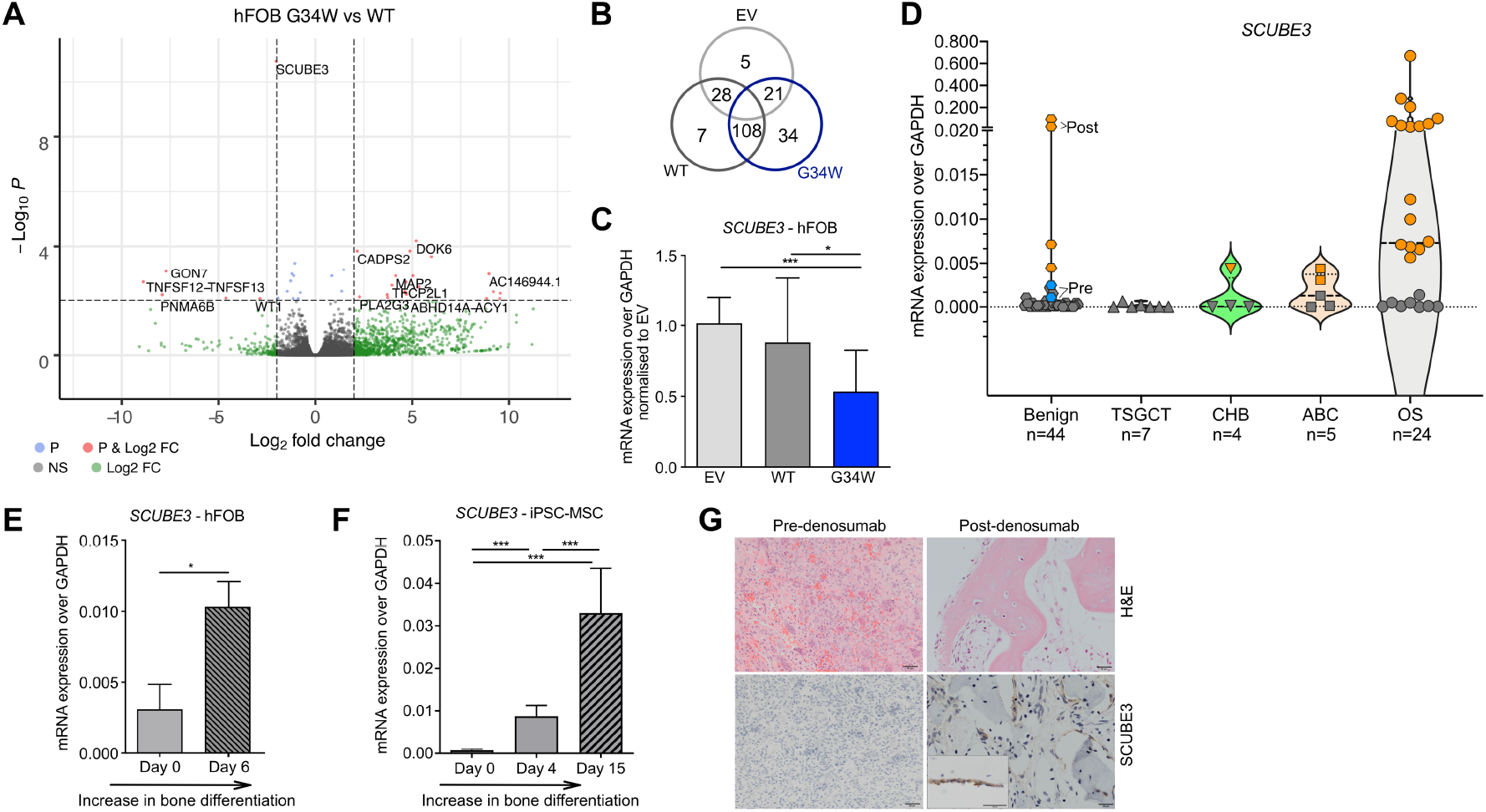
H3.3^G34W^ hFOB cells and GCTs have low levels of *SCUBE3*. **A**. Volcano plot showing the relationship between the mean difference in gene expression by RNA-seq; p < 0.001 (i.e. -log10P >3) and log fold Change >2. Total 35,639 variables. P is the Independent Hypothesis Weighting (IHW)-adjusted p-value. **B**. Venn diagram showing the number of differentially expressed genes in hFOB transfectants by RNA-seq. **C**. Expression of *SCUBE3* by qPCR in hFOBs. 6 replicates, 4 independent infections. **D**. Expression of *SCUBE3* by qPCR in GCT and GCT-mimics: benign GCT, TSGCT-tenosynovial giant cell tumour, CHB-chondroblastoma, ABC-aneurysmal bone cyst and OS-osteosarcoma. Grey: osteoclast-rich samples. Orange: bone-forming samples. In benign GCT: top two samples are post-denosumab treated from 2 patients (Post, orange), and their corresponding osteoclast-rich pre-denosumab samples (Pre, light blue). Dashed lines: median and quartiles. **E-F**. Expression of *SCUBE3* by qPCR in (E) H3.3^WT^ hFOB undifferentiated and after 6 days of differentiation (6 replicates, 2 experiments) and in (F) H3.3^WT^ iPSC-derived-MSCs during osteoblast maturation (6 replicates, 2 experiments). **G**. H&E-stained sections showing SCUBE3 in mature bone lining cells of post-denosumab GCT patients’ samples; 20X magnification. Data are mean+/-SD. C,F: 1-way ANOVA. E: unpaired Student’s t-test.

### H3.3^G34W^ stimulates osteoclast recruitment

We next tested if the H3.3^G34W^ mutation in osteoprogenitors accounts for the prominent osteoclast population in GCT. Using an *in vitro* human osteoclastogenesis assay we showed that the conditioned medium (CM) from differentiated (bone-forming) H3.3^G34W^-hFOBs significantly increased the number of osteoclasts and particularly those with more than 10 nuclei, albeit the results were variable due to donor-to-donor variability (**Figure 1G-I**). CM from differentiated H3.3^G34W^-stromal cells from iPSC-MSCs also induced a significant increase in large osteoclasts (**Figure 1J-L**). This supports the concept that the driver mutation in stromal/osteoprogenitor cells is responsible for the osteoclast-rich phenotype of the GCT.

### H3.3^G34W^ affects chromatin and transcription of osteoprogenitors leading to downregulation of *SCUBE3*

We next sought the molecule(s) underlying the osteoclast-inducing effect by performing bulk RNA-sequencing (RNA-seq) of H3.3^G34W^ hFOB and control lines (**Figure 2A-B, Supplementary Figure 3A-B** and **Supplementary Data 1**).

**Figure 3.**
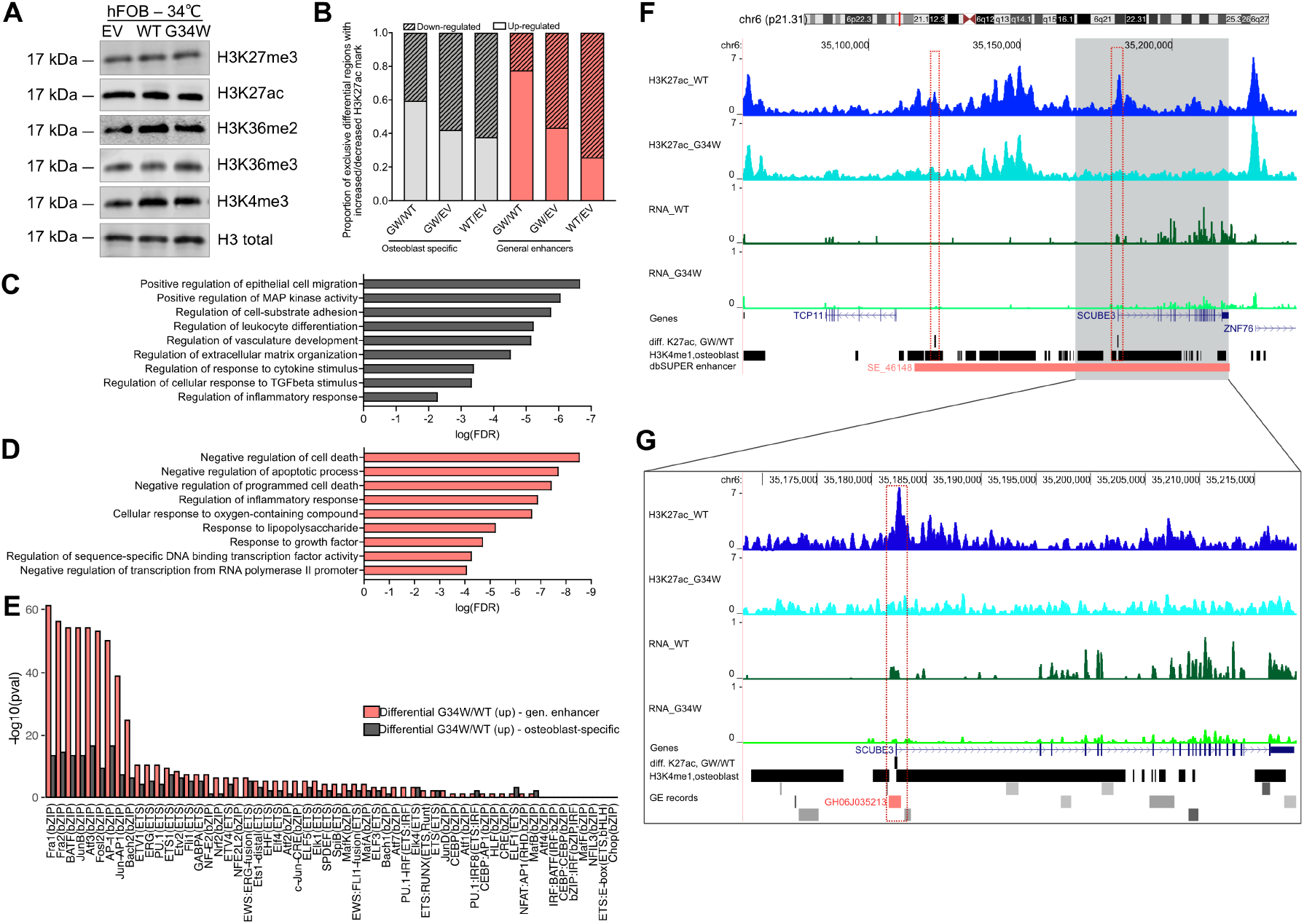
H3.3^G34W^ alters histone levels at enhancers of genes critical for osteoblasts, including at *SCUBE3* which regulates osteoclast formation. **A**. Total levels of histone marks on histone preparations by western blot of undifferentiated hFOB transfectants. **B**. Proportion of exclusive differential peaks showing increased (up-regulated) or decreased (down-regulated) H3K27ac marks from pairwise comparison among hFOBs transfectants, intersected with osteoblast-specific H3K4me1 genomic regions and general human enhancers. **C-D**. Gene Ontology (GO) analysis of genes neighbouring differential H3K27ac peaks up-regulated exclusively in G34WvsWT intersected with (C) osteoblast-specific H3K4me1 regions or with (D) human general enhancers. **E**. Significance of AP-1 and Ets TFs-motifs in differential H3K27ac peaks exclusively up-regulated in G34WvsWT, intersected with osteoblast-specific H3K4me1 regions (grey) and general human enhancers (coral). **F**. H3K27ac modifications (top pair of tracks) at the *SCUBE3* locus and RNA expression (3rd and 4th tracks) in H3.3^WT^ and H3.3^G34W^-hFOB. Differential peaks GWvsWT are shown in black. The super-enhancer SE_46148 (reported in dbSUPER database) is in red. Tracks show one representative replicate. **G**. As (F) but with an expanded view of the *SCUBE3* gene. The enhancer GH06J035213 (reported in GE database) is in red.

Notably, we found that H3.3^G34W^ had no impact on the expression of *RANKL* and *osteoprotegerin*, as previously described (10), and other critical cytokines involved in osteoclast formation (12) (**Supplementary Figure 3B-C** and **Supplementary Data 1**). Instead, the *Signal Peptide CUB Domain And EGF Like Domain Containing 3 (SCUBE3)*, a secreted member of the transforming growth factor beta (TGFβ) family, was the most significantly downregulated gene in H3.3^G34W^ compared to H3.3^WT^ cells (**Figure 2A,C**).

SCUBE3, which acts as an endogenous ligand for TGFβ-receptor 2 (13), is a valid candidate for regulating osteoclast recruitment: it is expressed by osteoblasts (14) (**Supplementary Figure 3D**), its inactivation results in a syndrome characterised by impaired ossification in humans and mouse models (15,16), and it is linked to Paget’s disease of bone characterised by numerous large osteoclasts (14). Consistent with our *in vitro* results, *SCUBE3* mRNA was virtually absent in GCT samples and in other osteoclast-rich tumours (**Figure 2D**). Instead, *SCUBE3* mRNA was increased in post-denosumab-treated GCTs and also in bone-forming osteoclast-poor osteosarcomas (**Figure 2D**). These data suggest an inverse correlation between osteoprogenitor *SCUBE3* expression and osteoclast activity across several bone tumour types, an effect potentially related to the state of maturation of bone cells. This is supported by higher levels of *SCUBE3* in osteoblasts undergoing differentiation *in vitro* (**Figure 2E-F**) and by the localisation of SCUBE3 to bone-lining cells in human samples of mature bone (**Figure 2G**).

Given the role of oncohistones in disrupting physiological chromatin states, we next investigated whether H3.3^G34W^ caused alterations in histone marks at the *SCUBE3* locus and other genomic regions. Unlike previous reports on the effect of other oncohistones including H3.3^K27M^ and H3.3^K36M^ (17–19), western blot analyses did not reveal global alterations in H3K27ac and other histone marks in H3.3^G34W^ hFOB (**Figure 3A** and **Supplementary Figure 4A**).

**Figure 4.**
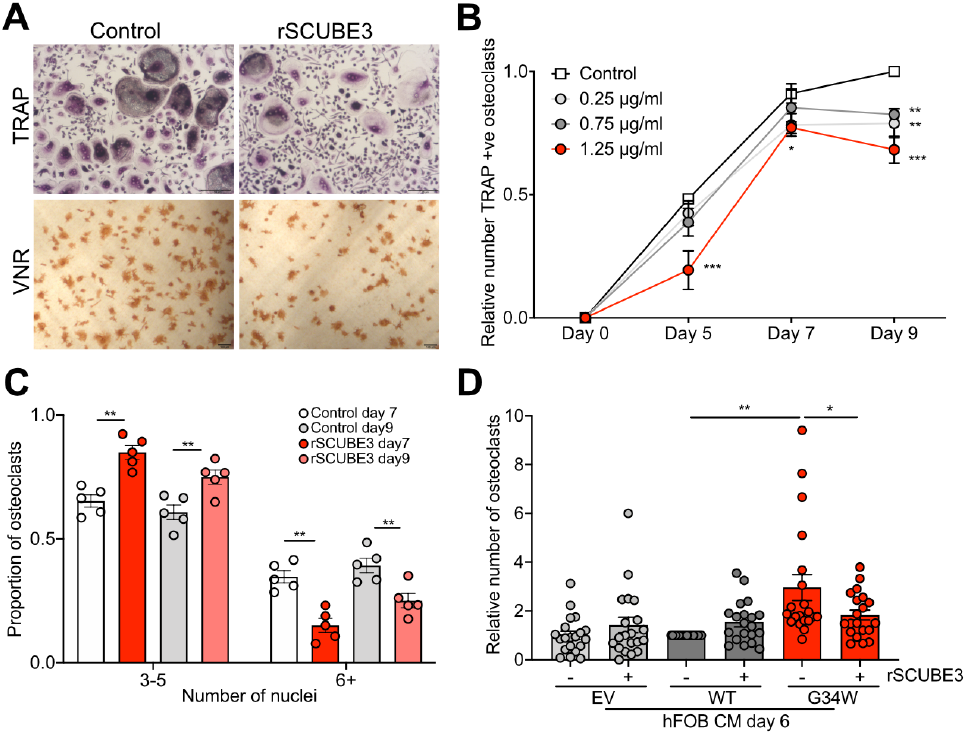
*SCUBE3* regulates osteoclast formation. **A**. Representative photomicrographs of rSCUBE3-treated osteoclasts on day 9 of differentiation showing inhibition of TRAP and vitronectin receptor (VNR) expression. **B-C**. Dose-dependent inhibition of the number of (I) total TRAP-positive osteoclasts and (J) osteoclast fusion by rSCUBE3; 5 preparations. **D**. rSCUBE3 inhibits the number of osteoclasts induced by CM only from differentiated H3.3^G34W^ hFOBs; 20 preparations. Data are mean+/-SEM. I-J: 2-way RM ANOVA. K: 1-way ANOVA.

Reasoning that H3.3^G34W^ may affect cell identity/fate by modifying chromatin states at enhancers, similar to H3.3^K27M^-mutated gliomas (20), we profiled hFOBs by ChIP-sequencing normalised with an exogenous reference genome for the enhancer mark H3K27ac (**Supplementary Figure 4B-C and Supplementary Data 2-3**). A large number of regions displayed differential H3K27ac enrichment, with an overall increase in H3.3^G34W^ (GW) compared to WT and EV (**Figure 3B**). The majority of differentially enriched peaks (GW/WT) overlapped with general human enhancers (12%) and mostly with reported osteoblast-specific H3K4me1 genomic regions (71%) (**Supplementary Figure 4D-F**) which appeared skewed towards promoter regions (**Supplementary Figure 4G**).

We also profiled the distribution of H3K36me3 mark, as it is altered by H3.3^G34R^ in brain cancer (19,21) (**Supplementary Figure 4H-L** and **Supplementary Data 4-5**). At regions of differential enrichment, H3K36me3 was mostly gained in the presence of H3.3^G34W^ (**Supplementary Figure 4M-N**), in close proximity to genes involved in regulation of cell shape and bone metabolism (**Supplementary Figure 5A-C**). Similarly, H3K27ac differentially enriched regions were neighbouring genes involved in cell migration, extracellular matrix organisation, response to cytokine stimulus as well as inflammation-related pathways and negative regulators of cell death (**Figure 3C-D** and **Supplementary Figure 5D-I)**. Transcription factor (TF) motif analysis revealed enrichment of AP-1/AP1-related motifs and Ets/Ets-like motifs (**Figure 3E** and **Supplementary Figure 5J-K**) suggesting a role for AP1-related and Ets-like TFs, known to control bone homeostasis and bone tumour development (22,23). Overall, these findings show that H3.3^G34W^ alters the chromatin at genes involved in osteoblast biology, which could explain the histological phenotype of GCTs (5,10).

**Figure 5.**
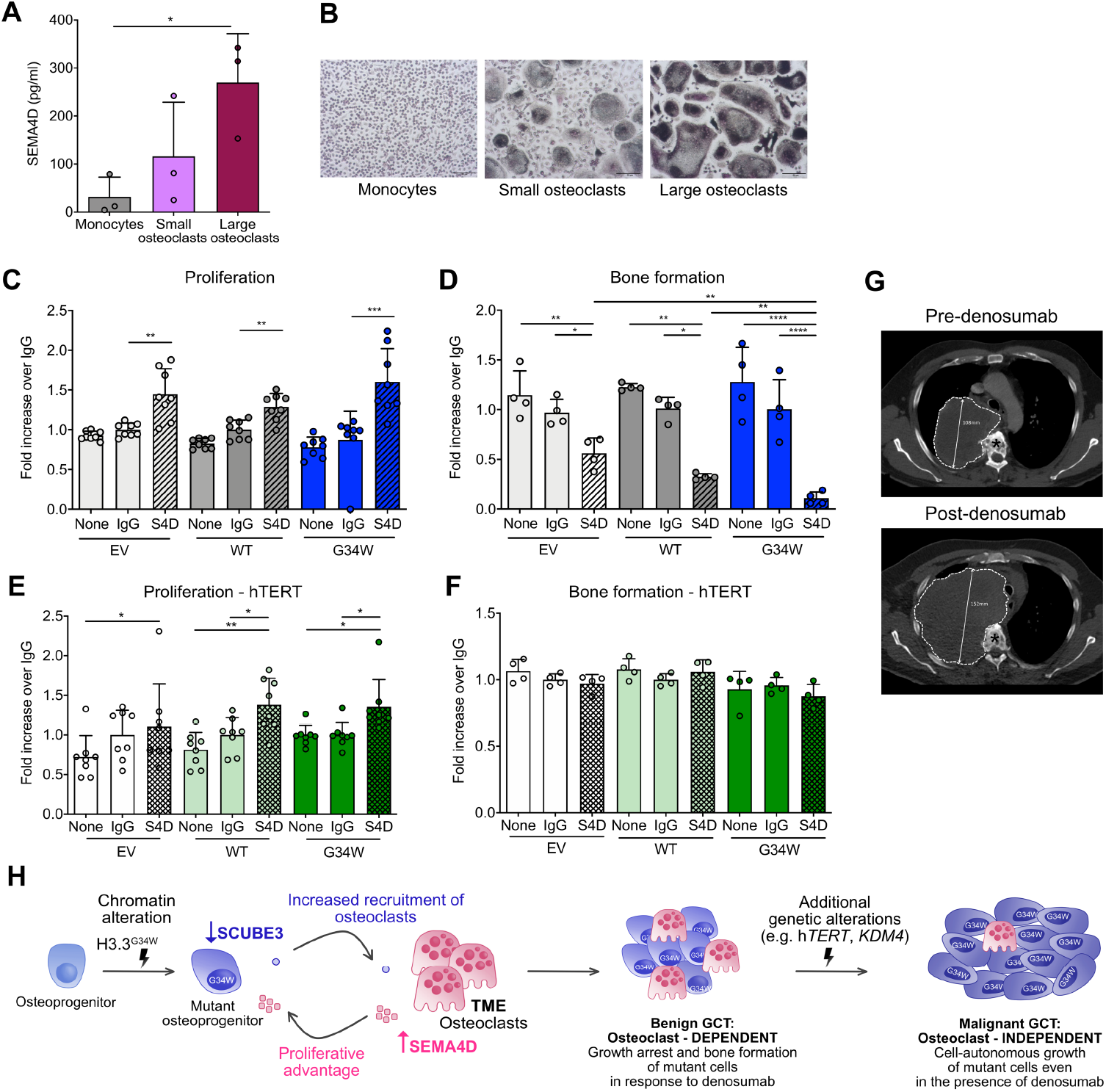
Benign and malignant GCTs respond differently to the TME. **A**. ELISA for SEMA4D on the supernatant of human monocytes and small/large osteoclasts after 9 days of differentiation; 3 preparations. **B**. Representative TRAP images of cells in (A). **C**. Proliferation of hFOB transfectants in the presence of rSEMA4D, IgG control or mineralization medium only (none) by Presto Blue assay after 7 days of proliferation at 34°C; 8 replicates. **D**. Osteoimage assay of hFOB transfectants on day 6 of differentiation; 4 replicates. **E**. Proliferation of hTERT-hFOB transfectants by Presto Blue assay after 7 days of proliferation at 34°C; 8 replicates. **F**. Osteoimage assay of hTERT-hFOBs on day 6 of differentiation; 4 replicates. **G**. Malignant GCT treated with denosumab: axial CT at T4-T5 vertebral level pre-and 3 months post-denosumab. In contrast to a conventional GCT, this tumour has grown from 108mm to 152mm (dotted outline) and has not induced mineralisation. Asterisk, vertebral body. **H**. Proposed schema for GCT evolution. H3.3^G34W^-mutant osteoprogenitors express reduced levels of SCUBE3 resulting in increased formation of large osteoclasts, which secrete high levels of SEMA4D that block differentiation and promote proliferation of H3.3^G34W^-osteoprogenitors. The transition from benign to malignant GCT requires acquisition of at least one additional genomic alteration and malignant cells display cell-autonomous growth. Data are mean+/-SD. A,C-F: 1-way ANOVA.

On inspection of the *SCUBE3* locus, we found that H3K36me3 distribution was not affected by H3.3^G34W^ (**Supplementary Figure 6A-B)**, whereas H3K27ac displayed differential enrichment at the gene regulatory region. H3K27ac was significantly reduced in H3.3^G34W^ compared to H3.3^WT^ cells (**Figure 3F-G**) in a region that overlaps with an osteoblast-specific super-enhancer, consistent with the observed reduction in *SCUBE3* mRNA expression. Interestingly, H3K27ac was also altered (positively or negatively) at regions in the proximity of a large set of secreted factors, including members of the TGFβ pathway, to which SCUBE3 belongs, suggesting a wider role of H3.3^G34W^ in epigenetic regulation of the osteoblast secretome (**Supplementary Figure 6C**).

### Reduced SCUBE3 expression contributes to increased osteoclast formation

Next, we assessed whether downregulation of *SCUBE3* contributed to the pro-osteoclastogenic effect of H3.3^G34W^-derived CM. Recombinant SCUBE3 (rSCUBE3) reduced the overall number and size of osteoclasts generated *in vitro* (**Figure 4A-C** and **Supplementary Figure 7A-C**), a finding supported by the altered expression of *OCSTAMP* and *DCSTAMP*, genes involved in osteoclast fusion (**Supplementary Figure 7D-E**), without inducing cell death (**Supplementary Figure 7F-H**). rSCUBE3 also counteracted the osteoclastogenic effect of H3.3^G34W^-hFOB-CM (**Figure 4D**), suggesting that the abundant giant osteoclasts in GCTs are the consequence of a paracrine effect driven by the H3.3^G34W^-osteoprogenitors.

### Osteoclast-secreted SEMA4D provides the growth advantage to GCT

Next, we speculated on the nature of the osteoclast-derived factor that provides the growth advantage to the tumour cells. We reasoned that Semaphorin 4D (SEMA4D), one of the few osteoclast-produced molecules known to suppress bone formation (12,24,25), which is expressed by osteoclasts in GCT (**Supplementary Figure 7I**), represents a good candidate. Primary cultures of ‘large’ osteoclasts produced greater amounts of secreted SEMA4D compared with smaller osteoclasts (**Figure 5A-B)**. Treatment of hFOBs with recombinant SEMA4D (rSEMA4D) promoted proliferation equally in the three transfectants (**Figure 5C**) whereas rSEMA4D exerted a more pronounced reduction in bone formation on the H3.3^G34W^-hFOBs (**Figure 5D**). Therefore, SEMA4D could confer the growth advantage to mutant cells in GCT by simultaneously enhancing their proliferation and blocking differentiation.

### Malignant GCTs do not rely on osteoclasts for growth

We next asked if the osteoclast-dependent growth advantage in conventional GCTs also occurs in malignant GCTs. We investigated the effect of the overexpression of *hTERT*, a common driver in malignant GCTs (3), on hFOB transfectants. Irrespective of the H3.3 mutation status, *hTERT* induced a proliferative advantage (**Supplementary Fig 7J-K**) but did not alter differentiation (**Supplementary Figure 7L**). Treatment of hTERT-hFOBs with rSEMA4D increased proliferation of the three transfectants (**Figure 5E**), although the effect was minor compared to that induced by hTERT (**Supplementary Fig 7K)**. However, rSEMA4D failed to block maturation of hTERT-cells (**Figure 5F**), a finding consistent with the common occurrence of bone formation in malignant GCTs (26,27). These results suggest that the histone mutation is not required for sustaining growth after malignant transformation (3,28) and that denosumab treatment would not curtail malignant disease. This concept is supported by the observation that denosumab treatment was ineffective in a patient with a malignant GCT harbouring a *hTERT* mutation (**Figure 5G** and **Supplementary Figure 7M-O**).

## Discussion

The TME is known to play an important role in neoplasia (36). Here we show that the main genetic driver event, the H3.3^G34W^ mutation, in cells of the osteoblast lineage in benign GCTs exerts its growth-promoting effect in a non-cell autonomous manner through osteoclasts in the TME (**Schema** in **Figure 5H**). Our evidence is supported by the clinical effect of denosumab, which results in growth arrest and maturation of the tumour cells into bone-forming cells. However, to date, the osteoclast-produced molecule(s) that promotes growth of the mutant osteoprogenitors has never been identified. Using human *in vitro* models, we show here that SEMA4D provides a proliferative advantage to mutant osteoprogenitors. Our data also provide evidence that malignant GCTs would not benefit from treatment with denosumab: this assertion is made on our experimental evidence that mutant osteoprogenitors with a hTERT-mutated phenotype, characteristic of malignant GCTs (3), largely loose their dependency on osteoclasts for their proliferative advantage. Furthermore, our finding that SEMA4D does not block osteoblast differentiation in the presence of hTERT over-expression explains why malignant transformation of GCT can result in an osteosarcomatous histological phenotype (33).

Mechanistically, we find that H3.3^G34W^ in the absence of a proliferative advantage has a direct effect on osteoblast precursors by altering the expression of genes involved in extracellular matrix organisation and by inducing a block in osteogenic differentiation, a finding consistent with the histological phenotype of GCTs as also recently suggested by others (5,18). We demonstrate that H3.3^G34W^ modulates the profile of enhancers in the proximity of a large set of osteoblast-secreted factors, including *SCUBE3*, a molecule involved in the physiological bone growth and ossification process (24): we provide evidence that reduction in expression of SCUBE3 is involved in the conspicuous osteoclast population in GCTs.

*H3-3A* and -*3B* are paralogous genes that transcribe the same product. Mutations in these genes occur in bone and paediatric brain tumours (2,11) but the associated tumour type is largely specific for both the gene and mutation (37). Our study shows that the H3.3^G34W^ mutation is associated with alterations of the epigenetic landscape of the mutant osteoprogenitors which notably comprise osteoblast-specific enhancers, including that of *SCUBE3*, thereby implicating this mutation in the pathogenesis of GCT specifically and not in other tumours. However, it remains to be established whether H3.3^G34W^ is able to modulate H3K27ac levels directly via some yet-to-be determined molecular mechanisms, or if the observed changes in H3K27ac are more a reflection of the cell of origin and/or differentiation stage at which this mutation arises. These two hypotheses are not mutually exclusive, as the mutant histone could be able to promote enhancer changes associated with earlier developmental stages, thus stimulating dedifferentiation of a more mature osteoblastic cell.

Collectively, our results provide evidence at a molecular level for a bidirectional communication between tumour cells and osteoclasts in benign GCT, a dependency which is largely lost on malignant transformation. Moreover, our work provides a starting point for the identification of similar processes in other mesenchymal neoplasms which will lead to understanding the role of critical molecules, such as SCUBE3, in health and disease.

## Materials and methods

### Vector construction

pHIV-dTomato lentiviral vector (a gift from Bryan Welm, Addgene plasmid # 21374) was used to transduce hFOB and iPSC. H3.3^WT^ and H3.3^G34W^ vectors were produced by linearising pHIV-dTomato using EcoRI to insert cDNA encoding C-terminal HA-tagged Drosophila *His3*.*3A* (a highly conserved ortholog of mammalian *H3-3A*) followed by the IRES and the dTomato sequences: GGA encoding for Glycine at position 34 was changed to TGG encoding for Tryptophan at position 34. The empty vector (EV) was used as a control. Lentiviral particles were produced as described previously (29). hTERT was overexpressed in hFOB transfectants using the pCWX-UBI-hTert-PGK-BSD lentiviral vector (a gift from Patrick Salmon, Addgene plasmid #114316).

### Cell culture

Cell lines were expanded and differentiated according to the following protocols and cell authentication was regularly performed by Short Tandem Repeat fingerprinting (Culture Collections, Public Health England, UK) (**Supplementary Table 2**). Regular testing was also performed to ensure that the cell lines were mycoplasma-free using the EZ-PCR Mycoplasma Test Kit (K1-0210, Geneflow, Lichfield, Staffordshire, UK). <u>hFOB</u>. hFOB1.19 ATCC® (CRL-11372™, ATCC, Manassas, VA, USA), a human fetal osteoblastic cell line immortalised using temperature-sensitive SV40 large T antigen which proliferates at the permissive temperature of 34°C and undergoes osteoblastic differentiation at 39°C (30), was grown in Dulbecco’s Modified Eagle Medium/Nutrient Mixture F-12 (21041025, Gibco Life Technologies, Thermo Fisher Scientific, Loughborough, Leicestershire, UK) supplemented with Fetal Bovine Serum (FBS) (F9665, Sigma Aldrich, St. Louis, MO, USA) to a final concentration of 10%, hereafter referred to as normal medium as described in the original publication (30). This medium was supplemented with Geneticin® Selective Antibiotic (G418 Sulfate, 50 mg/mL, 10131035, Sigma Aldrich), which was added during maintenance culture (at 34°C) but omitted when performing the osteogenic assays.

hFOB were infected with lentiviral particles an MOI of 10, incubated over night at 34°C in the presence of the virus which was removed the next morning. The cells were expanded and FACS-sorted for dTomato. hFOB experiments were performed using cells derived from 4 independent infections. All experiments were performed using cells >90% dTomato positive. To generate h*TERT*-expressing cells, hFOB-EV, -WT and –G34W cells were infected with lentiviral particles produced as described above, followed by antibiotic selection using Blasticidin S HCl (Thermo Scientific, A1113903, final concentration 5µg/ml) and the surviving cells expanded for at least 3 passages before being used for functional assays.

To differentiate hFOB to mature osteoblasts (**Supplementary Figure 1**), cells were seeded at a density of 0.4×10^6^ cells/well in 24 multiwell plates coated with collagen (Collagen, C3867, Sigma Aldrich), grown over night at 34°C, then moved to 39°C either in normal medium or in the presence of Osteoblast Mineralisation Medium (C-27020, Promocell, Heidelberg, Germany) and grown for 6 days or until signs of mineralisation were evident (up to 12 days). Conditioned medium (CM) of hFOBs grown in the absence of Geneticin was collected from cells seeded at a density of 0.4×10^6^ cells/well and grown overnight at 34°C (day 0) or after 6 days at 39°C and grown in mineralisation medium. Cell debris were removed by spinning at 2500 rpm for 5 minutes and CM was kept at -80°C until needed.

#### Human iPSC-derived MSC

The viral-integration-free human iPSC line generated using cord blood-derived from CD34+ progenitors was obtained from Gibco™/Thermo Scientific (Cat. A18945) and grown in serum-free culture conditions according to the manufacturer’s instructions. Briefly, cells were cultured and expanded in 6-well plates coated with Geltrex™ LDEV-Free Reduced Growth Factor Basement Membrane Matrix (A1413202, Gibco Life Technologies) diluted 1:100 in DMEM (31966021, Gibco Life Technologies). Cells were maintained in Essential 8™ Flex Medium Kit (A2858501, Gibco Life Technologies) and passaged twice per week on reaching 80-90% confluency, using 0.5mM EDTA in Dulbecco’s phosphate-buffered saline (14190250, Gibco Life Technologies); after which they were split (1:3 to 1:6) using trypsin. iPSC used in this study were between passages 40 and 75.

iPSC-derived MSC (**Supplementary Figure 2**) were obtained by seeding iPSCs at a density of 50000 cells/well in 12 well. 24 hours later, mesoderm differentiation was induced by adding Cardiomyocyte Differentiation medium A (A29209-01, Gibco Life Technologies) for 48 hours, after which iPSC-derived MSC were transduced. Cells were infected with lentiviral particles with an MOI of 15 using spinoculation (200-400g at 34°C for 30 minutes) and incubated overnight in the presence of virus, which was removed the following morning and replaced with fresh mineralisation medium or control MSC maintenance medium. iPSC-derived MSC cells were infected using fresh virus for each experiment and efficiency of infection was monitored by fluorescence microscopy. On the same day, osteoblast differentiation was induced by adding Mesenchymal Stem Cell Osteogenic Differentiation Medium (C-28013, Promocell) or Human Mesenchymal Stem Cell (hMSC) Osteogenic Differentiation Medium BulletKitTM (PT-3002, Lonza) according to manufacturer’s instructions. MSC used as control were passaged and maintained in MSCGM Mesenchymal Stem Cell Growth Medium BulletKitTM (PT-3001, Lonza). All steps were performed at 37°C.

CM of osteoblasts differentiated from iPSC-derived MSC was collected from cells seeded at a density of 25000/well in a 24 multiwell plate after 8 or 15 days of osteoblasts differentiation. Cell debris were removed by spinning at 2500rpm for 5 minutes and CM was kept at -80°C until needed.

#### Human osteoclasts

Human osteoclasts were generated as described previously(31): CD14+ monocytes were positively selected from the peripheral blood mononuclear cell component of leucocyte cones (NHS Blood and Transplant, UK) using CD14+ microbeads (130-050-201, Miltenyi Biotech, Surrey, UK). Monocytes were seeded onto dentine discs (elephant dentine; HM Revenue & Customs, Heathrow Airport, UK) or plastic dishes in α-MEM (without ribonucleosides/ deoxyribonucleosides; Lonza) containing 10% heat-inactivated FBS, 2 mM L-glutamine, 50 IU/ml penicillin and 50 μg/ml streptomycin sulphate. Osteoclastogenesis was induced by treatment with 25 ng/ml human M-CSF (216-MC, R&D Systems, Abingdon, UK) and 30 ng/ml RANKL (310-01, Peprotech, London, UK) every 3–4 days for 9 days. Small osteoclasts were generated for selected experiments using 3 ng/ml of RANKL, but otherwise were generated using 30 ng/ml of RANKL; monocytes were maintained in M-CSF in the absence of RANKL: CM for the ELISA was collected on day 9 of differentiation, 48 hours after the last medium change. Use of leucocyte cones for osteoclast differentiation was approved by the London -Fulham Research Ethics Committee (11/H0711/7).

### Flow cytometry Activated Cell Sorting (FACS)

After transduction, hFOB cells were expanded to reach at least 2×10^6^ cells, dissociated into single cells and sorted using a BD FACS Aria Fusion Cell Sorter ™ (Becton Dickinson, USA) running FACSDiva Software version 6. hFOB were bulk sorted to exclude DAPI+ dead cells and to select dTomato-expressing cells (aiming for >99.9% positive cells). Positivity for dTomato of sorted cells was checked over time in different passages by Flow Cytometry, on an LSR Fortessa™ (Becton Dickinson, USA) running FACSDiva Software version 6 with 10^4^ events recorded for each sample.

### Osteogenic assays

#### Alizarin red staining (ARS) and quantification

Mineralised osteoblasts were fixed and stained according to the ‘Detection of Calcium Deposits (Mineralisation)’ Promocell protocol (PromoCell GmbH website). Briefly, fresh 2% (40 mM) ARS solution was prepared by adding 2g of Alizarin (C.I.58005) to 100 mL of water and the pH was adjusted to 4.1-4.3 with 0.1% NH_4_OH. The solution was filtered and stored in the dark. Cells were gently washed with Phosphate Buffer Saline (PBS), fixed in neutral buffered formalin (10%) for 30 minutes, washed once with distilled water, incubated at room temperature in ARS solution for 45 minutes in the dark, washed 4 times with distilled water and kept in PBS. For the quantification, the stained cell monolayer was incubated at room temperature in 10% acetic acid for 30 minutes with shaking. Cells were collected using a cell scraper, vortexed for 30 seconds, heated at 85°C for 10 minutes, incubated on ice for 5 minutes and then centrifugated at 20,000g for 15 minutes. Supernatant was transferred to a new tube and 10% ammonium hydroxide added to neutralise the acid. pH was checked in a small aliquot to ensure it fell within the range 4.1-4.5. The absorbance was read at 405 nm with a plate reader.

#### OsteoImage assay

Cells were seeded at a density of 100000/well in collagen-coated 96 well plates, fixed after 12 days and stained according to the OsteoImage™ Mineralisation Assay Lonza kit (PA-1503, Lonza) protocol. Briefly, the cell monolayer was washed once in PBS, fixed in neutral buffered formalin (10%) for 30 minutes and rinsed in 1x OsteoImage™ Wash Buffer. OsteoImage™ staining reagent was added and incubated for 30 minutes at room temperature, protected from light. Cells were washed three times with washing buffer for 5 minutes. Fluorescence was read in a plate reader at excitation/emission wavelengths (492/520).

### Functional assays

#### Incucyte proliferation assay

hFOB were collected, counted and plated in TPP 96 well plates at 2500/well in at least 3 replicate wells per experiment. Cells were incubated at 34°C or at 39°C using an Incucyte Zoom® live cell imaging system (Essen BioScience, MI, USA). Images were taken every 2 hours for 7 days and confluency was calculated using the Incucyte software.

#### Colorimetric viability assay

Cell viability was measured using Presto Blue Cell Viability Reagent (Cat A13262, Thermo Fisher Scientific, Loughborough, UK) according to the manufacturer’s instructions.

#### Edu proliferation assay

iPSC-derived MSC cells were differentiated in osteoblasts or maintained in MSC medium for 2 days, collected, stained and analysed using the Click-iT™ EdU Cell Proliferation Kit for Imaging, Alexa Fluor™ 647 dye (Cat. C10340, Thermo Scientific) according to manufacturer’s instructions. Experiments were performed on an LSR Fortessa™ (Becton Dickinson, USA) running FACSDiva Software version 6 with 10^4^ events recorded for each sample.

#### Apoptosis assay

iPSC-derived MSC cells were differentiated into osteoblasts for 2 days and collected. hFOB were grown at 34°C until 70% confluency was reached and then collected. Apoptosis was determined by detecting phosphatidylserine using the APC-Annexin-V Apoptosis Detection Kit with PI (Biolegend, CA, USA). Briefly cells were harvested, washed once in PBS and 2×10^5^ cells resuspended in 250 µl of binding buffer containing 5 µL Annexin V-APC and 10 µl PI solution. Cells were incubated in the dark for 15 minutes before being analysed. Each assay was repeated 3 times, each with 3 replicates. Experiments were performed on an LSR Fortessa™ (Becton Dickinson, USA) running FACSDiva Software version 6 with 10^4^ events recorded for each sample.

#### Wound healing assay

A monolayer scratch assay was performed by seeding 30000 cells/well in 24 well ImageLock plates (Essen Instruments, Cod. 4365) and incubating them for 3-4 days. hFOB were grown on plates coated with Collagen I solution (Sigma) at 34°C. When cells were 95% confluent, wounds were created using the EssenBio wound maker and plates were scanned for 48h using the Incucyte™ FLR live cell imaging system (Essen BioScience, MI, USA). The system measures scratch closure in real time and automatically calculates the relative wound density within the initially empty area over a time course.

### Osteoclast formation, activity and survival assays

Tartrate-resistant acid phosphatase (TRAP) and the vitronectin receptor (CD51/61, VNR) are osteoclast markers used for the visualisation of mature osteoclasts (33). TRAP staining was performed on formalin-fixed cells using naphthol AS-BI phosphate as a substrate, with reaction of the product with Fast Violet B salt. Multinucleated cells containing three or more nuclei were considered osteoclasts. VNR was detected on cells fixed in cold methanol by CD51/61 immunocytochemistry (clone 23C6, 1:400; Bio-Rad, Oxford, UK). Resorption tracks on dentine discs were visualised by staining with 0.5% toluidine blue under reflected light. The dentine slices were photographed, resorption tracks highlighted, and the resorbed area quantified using ImageJ. Terminal deoxynucleotidyl transferase dUTP nick end labelling (TUNEL) staining was performed using the In Situ Cell Death Detection Kit, POD (Sigma).

### Recombinant proteins and conditioned medium (CM) treatments

Effects of rSCUBE3 were investigated by treatment of osteoclast cultures with 0.25-1.25 µg/ml recombinant human rSCUBE3 protein (Cat # 7730-SC, R&D Systems, Abingdon, UK) in the presence of RANKL and M-CSF; all the reagents were replaced at each media change for osteoclasts. For control experiments, rSCUBE3 was denatured at 95°C for 30 minutes. Effects of osteoblast CM on osteoclasts were investigated by adding 10% CM generated from EV, WT or G34W hFOB or MSC-derived osteoblasts each time the medium was changed. Effect of rSEMA4D was investigated by treating osteoblasts with recombinant Human Semaphorin 4D, Fc Tag 15μg/ml (CDO-H5257, Acro Biosystem Newark, USA) or Recombinant Human IgG1 Fc 3μg/ml (110-HG, R&D Systems, Minneapolis, MN, USA) as control. hFOB were seeded in collagen-coated 96 well plates (TPP) in 4 wells per genotype per condition at a density of 1×10^5^ cells/ well and kept at 34°C for 24 hours. For proliferation assay (colorimetric assay), the cells were kept in hFOB medium at 34°C for 7 days. For bone formation assay the medium was replaced with Mineralisation medium (Promocell) and the plates moved to 39°C for 12 days. Spent medium was replaced with fresh medium including fresh rSEMA4D/IgG every 4 days.

### RNA extraction and qPCR

Total RNA was extracted using miRNeasy Mini Kit (217004, Qiagen, Manchester, Lancashire, UK). Quantitative real-time PCR (qPCR) was performed as previously described (34). FFPE and fresh frozen GCT tissue samples were processed for RNA extraction as described in Cottone at al. (35). Primers used for qPCR are listed in **Supplementary Table 3**.

### Western blot and ELISAs

#### Western blots

were performed as described in Scheipl et al (34). Histone extraction was prepared according to Abcam Histone Extraction protocol (Abcam website). Antibodies used for western blot are listed in **Supplementary Table 4**.

#### ELISA

Protein quantification of CM from osteoclast cultures was performed using the Human SEMA4D (Semaphorin-4D) ELISA Kit (EH2196) (Wuhan Fine Biotech Co., China) according to manufacturer’s instructions.

### Immunohistochemistry (IHC) and Immunofluorescence (IF)

#### GCT samples

Tumour diagnoses were made using the WHO classification (WHO, 2020). Formalin-fixed paraffin-embedded (FFPE) samples were obtained from the archive of the Royal National Orthopaedic Hospital.

#### IHC

Hematoxylin and Eosin (H&E) staining and IHC were performed as described previously in (26) using the antibodies listed in **Supplementary Table 4**.

#### IF/Immunocytochemistry on iPSC

To confirm the expression of pluripotency markers in iPSC, the Pluripotent Stem Cell 4-Marker Immunocytochemistry Kit (Invitrogen™, Cat. A24881) was used according to manufacturer’s instructions. Images were acquired using an Axio Observer Z1 With Apotome.

#### Immunofluorescence

for Ki67 was performed as described previously (29) using the antibody listed in **Supplementary Table 4**, on transduced MSC after differentiated to osteoblasts for 3 days. Quantification of Ki67 positive nuclei was performed analysing 10 images (20X magnification) per condition.

### RNA sequencing

hFOB grown at 34°C for 15 days after transduction were collected in duplicate. Cells were lysed in Trizol and total RNA extracted using the Direct-zol kit (Zymo Research, CA, USA) including an on-column DNA digest. Poly(A) RNA was selected using the NEBNext Poly(A) mRNA Magnetic Isolation Module (New England Biolabs) and a first strand library prepared using NEBNext Ultra Directional RNA Library Prep Kit (New England Biolabs) and sequenced on a HiSeq 2500 (Illumina).

#### cDNA library construction and Illumina RNA-Seq

The cDNA libraries were constructed and sequenced by Source Bioscience, UK in accordance with the Illumina TruSeq RNA sample preparation guide v2 for Illumina paired-end multiplexed sequencing. In brief, the poly-A-mRNA in the extracted total RNA samples was purified using Illumina poly-T oligo-attached magnetic beads in two rounds of purification steps according to the manufacturer’s instruction. During the second step of poly-A RNA elution, the mRNA was fragmented and primed with random hexamers for cDNA synthesis. The first strand cDNA was synthesised from fragmented mRNA using reverse transcriptase and random primers. In a subsequent step, the RNA template was removed and a replacement was synthesised to construct double-stranded cDNA. After double-stranded cDNA synthesis, ends were repaired and an A-base was added to the blunt end fragments. Thereafter, Illumina indexing adapters were ligated according to the standard protocol for pooling of samples prior to sequencing and for subsequent identification of pooled samples in downstream analysis. The cDNA fragments that have adapter molecules on both ends were subjected to 15 rounds of PCR amplification. The concentration and size distribution of the synthesised cDNA libraries were confirmed using an Agilent BioAnalyzer 2100. The successfully amplified and indexed libraries were pooled and diluted to 10 nM prior to sequencing (two samples per lane). The molarity and size distribution were confirmed using an Agilent BioAnalyzer 2100. Finally, pooled samples were loaded at a concentration of 8 pM into each lane of an Illumina HiSeq 2000 flow cell v3 and sequenced with 100 bp paired-end reads.

#### RNA-seq data processing

The quality of the RNA-Seq data was examined using the package FastQC (http://www.bioinformatics.babraham.ac.uk/projects/fastqc/). RNAseq expression count estimates were made using kallisto software (37) together with the Ensembl GRCh38 (v99) transcript models. RNAseq count data were then imported using tximport (38) to the DESeq2 R package (39) for pre-processing, normalisation and statistical analysis. Multiple hypothesis adjustments used the independent hypothesis weighting method (IHW) (40). Principal component analysis was performed on rlog transformed expression values (39).

### Chromatin Immuno Precipitation-sequencing normalised with an exogenous reference genome (ChIP-Rx)-sequencing

hFOB grown at 34°C after transduction were collected in triplicate and washed once in cold PBS-5nM Na butyrate.

#### ChIP-Rx

A fixed ratio of *Drosophila* S2 cells (20% of hFOB cells) was spiked in prior to fixation to allow for exogenous normalisation. Cells were then fixed in 1% formaldehyde for 8 minutes prior to quenching with excess glycine. Fixed cells were resuspended on ice in wash buffer1 (10 mM Hepes pH7.5, 10mM EDTA, 0.5 mM EGTA, 0.75% Triton X-100, all reagents from Sigma-Aldrich) for 5 minutes, rotating at 4°C, centrifuged, and resuspended in wash buffer2 (10 mM Hepes pH 7.5, 200mM NaCl, 1mM EDTA, 0.5 mM EGTA, 0, all from Sigma-Aldrich) for 5 minutes rotating at 4°C. Samples were then diluted with Lysis Buffer (150 mM Na-HCL, 25 mM Tris pH 7.5, 5 mM EDTA, 1% Triton X-100, 0.5% Deoxycholate, 0.2% SDS, all from Sigma-Aldrich) and sonicated on a Bioruptor Pico (Diagenode, Belgium) for 6-10 cycles of 30 seconds on/30 seconds off. After sonication, Triton was added to a final concentration of 1%. Sonication efficiency was checked by running a sample of de-crosslinked material on a 2 % agarose gel. Equal amount of sonicated chromatin was incubated overnight rotating at 4ºC with the antibodies reported in **Supplementary Table 4**. Samples were incubated with protein A/G magnetic beads (Invitrogen) at 4ºC for 3 hours. Beads were washed sequentially with buffer 1 (50 mM Tris, 500 mM NaCl, 1 mM EDTA, 1 % Triton X-100, 0.1 % sodium deoxycholate, 0.1 % SDS, all from Sigma-Aldrich) three times, buffer 2 (20 mM Tris, 1 mM EDTA, 250 mM LiCl, 0.5% NP-40, 0.5% sodium deoxycholate, all from Sigma-Aldrich) three times and twice with TE buffer + 50 mM NaCl. DNA was eluted in buffer containing 50 mM Tris, 10 mM EDTA and 1% SDS before treatment with proteinase K and RNAse-A (both Thermo Fisher Scientific). DNA was purified with Qiaquick PCR purification Kit (Qiagen).

#### ChIP-Rx libraries construction and sequencing

Libraries were prepared using the NEBNext Ultra 2 DNA Library Preparation Kit (New England Biolabs, MA, USA) with AMPure® XP Beads (Beckman Coulter) and sequenced on a Nextseq500 (Illumina, CA, USA).

#### ChIP-Rx data processing

Raw data processing and alignment: low quality bases were trimmed, and adaptors were removed by Trimgalore with default parameters. Processed reads were aligned to hg19 and dm6 with “--no-unal” parameter. Alignment to hg19 was sorted, filtered to keep only normal chromosomes and indexed by Samtools. Initial scaling factors for each drosophila spiked-in IP samples were calculated following Niu, Liu, and Liu 2018. For each ChIP-type the initial scaling factors were adjusted by dividing initial scaling factors by the maximum initial scaling factor within the same ChIP antibody so that the maximum initial scaling factor was transformed to 1 and others accordingly. Filtered BAMs of IP samples were down-sampled by Picard with the adjusted scaling factor. BigWig files were generated from scaled BAMs by bamCoverage with binSize=1. Unique alignments from the filtered BAM files were subjected to peak calling by Homer using “histone” mode.

Integration of histone modification peaks from experimental replicates and H3K4me1/general enhancer: for G34W and EV samples (each has two replicates) the intersected peak regions from two replicates were identified; for WT samples (three replicates) peak regions that were intersected by at least 2 out of 3 replicates were identified. Integrated H3K27ac peak set for each genotype derived from the last step was intersected with either H3K4me1 peak or general enhancer. Only intersected regions long than 50 bp were kept.

Differential peak identification: scaled BAM files were transformed to BED format by “bam2bed” from BEDOPS. diffReps was used to find differential peaks between all replicates of any two genotypes with window=300 and meth=nb. Integration of differential H3K27ac peaks and H3K4me1/general enhancer: osteoblast H3K4me1 was downloaded from GEO (GSM733704 (42)) and general enhancer from FANTOM (https://fantom.gsc.riken.jp/5/datafiles/latest/extra/Enhancers/human_permissive_enhancers_phase_1_and_2.bed.gz). Both osteoblast-specific and general enhancer were intersected with differential peaks, and only intersected regions longer than 50bp were kept. Exclusive differential peak (for differential H3K27ac they have been intersected with H3K4me1 or general enhancer first) for any pairwise comparison from three genotypes (WT, G34W, EV) were defined as differential peaks that do not overlap with differential peaks from the other two pairwise comparisons. Up- and down-exclusive differential peaks are distinguished according to diffReps output. Venn diagrams showing the overlapping among differential peak sets were generated using ChIPseeker R package.

##### Functional analysis

GREAT was used for functional analysis on up (increased histone modification)-/down(decreased histone modification)-/both-exclusive differential peak sets with default setting. Results in GO-Biological process, mouse phenotype single knockout and mouse phenotype were downloaded, and only terms shown in the default GREAT result were used for assembling the heatmap. The color in the heatmap represents –log(hyperFDR).

Motif analysis: up (increased histone modification)-/down(decreased histone modification)-/both-exclusive differential peak sets were searched for enriched known motifs by Homer with size=300. The heatmap for enriched motifs was generated by first collecting the union set of top 20 enriched motifs from each peak list, and p-values for the union motif were extracted from the motif discovery result for each peak list. If the p-value was not found, it was assigned to 1.

Genomic feature distribution: ChIPseeker was used to generate barplot from BED files that derived from “Integration of histone modification peaks from experimental replicates and H3K4me1/general enhancer” section.

Tag intensity profile: computeMatrix and plotProfile from deeptools were used to generate the tag density profile over genic and TSS from the scaled BigWig files with binSize=10. Principle component analysis: PCA plot was generated from scaled BAM files (43).

Differential H3K27ac peaks in TGF-beta singling pathway: genes belonging to TGF-beta pathway were retrieved from MSigDB. The correspondence between differential H3K27ac peaks and nearby genes was identified by GREAT with default parameters. TGF signaling genes found in each differential peak list were assembled and represented as heatmap. Functional analysis for differential H3K27ac peaks between G34W and WT overlapped with general enhancer bearing ETS-related motifs: DNA sequence of differential H3K27ac peaks (GW/WT,up) overlapped with general enhancer was retrieved by “bedtools getfasta” and saved as FASTA file. MEME-formatted motif matrixes for selected ETS-related motifs (ETV4, ETV1, GABP1, EHF, ERG, PU.1, ELF5, Fli1, ETS1, ETV2) were downloaded from JASPAR and catenated into one single file. FASTA file and the catenated motif file were supplied to FIMO to identify ETS-bearing peak. ETS-bearing peaks were then fed to GREAT for functional analysis with default settings.

The global H3K36me3 level for each sample was calculated as in Pathania et al (44):

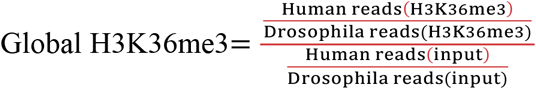

The following data are displayed in UCSC Genome Browser: Spike-in normalised IP signal from G34W and WT samples, BPM (Bins Per Million mapped reads) normalised BigWig for RNA-seq samples, differential H3K27ac peaks between G34W and WT, osteoblast H3K4me1 peaks from GEO (GSM733704(42)), super enhancer record for osteoblast from dbSUPER (http://asntech.org/dbsuper/index.php) and GeneHancer track from UCSC genome browser built-in.

## Data availability

High-throughput data (RNA-seq and ChIP-Rx) of hFOB cells have been deposited in the National Center for Biotechnology Information GEO database under GEO accession number GSE152942.

## Statistics

Statistical parameters including the exact value of n, precision measures (mean ± SD) and statistical significance are reported in the Figures and Figure Legends. In figures, asterisk denote statistical significance with the following symbols: *p ≤0.05, **p ≤0.01, ***p ≤0.001, ****p ≤0.0001. Continuous variables were compared via unpaired or paired t-test. Grouped data were analysed using 1-way or 2-way ANOVA with Tukey’s or Dunnett’s multiple comparison as a post hoc test. Data are always mean+/-SD. Statistical analysis was performed in GraphPad PRISM 8.0 (GraphPad Software, La Jolla, CA, USA).

## Study approval

Ethical approval for GCT samples was obtained from the Cambridgeshire 2 Research Ethics Service (reference 09/H0308/165) (HTA Licence 12198). Written informed consent was received from participants prior to inclusion in the study. All studies have been conducted according to Declaration of Helsinki principles.

**Supplementary information is available at Cell Death & Differetiation’s website.**

## Acknowledgments

We thank Diana Carvalho, UCL, for her contribution to preliminary experiments; Francesco Saverio Tedesco, UCL, for providing the iPSC cells; Teresa Sposito for help with iPSC cultures; Manuel Rodriguez Justo and Dominic Patel, UCL, for support with digital images; Ivana Bjedov, UCL, for providing S2 cells; Jenny Russ, PS group, for advice on NGS-related work; Dominique Heymann, Nantes University, for support. We also thank the Daniele Bano and Pierluigi Nicotera teams (DZNE Bonn) for support. We thank the UCL Cancer Institute Core Facilities, the DZNE Core Facilities, Platform foR SinglE Cell GenomIcS and Epigenomics (PRECISE, DZNE Bonn), the Biobank Team and the Research and Development Department at the RNOH, the healthcare workers and the patients for the generous donation of their material.

## Author contributions

Conceptualisation: AMF, PS. Study design: AMF, PS, LC, HK. Investigation: LC, LL, HK, SB, KH, JLS. Bioinformatic analysis: HML, SH, JH, APL. Clinical data and sample curation: AMF, FA, RT, SS, POD. Writing: AMF, LC, PS, AEG with input from other co-authors.

## Ethics statement

The study was performed in accordance with the Declaration of Helsinki.

