## Supplementary Material for "Aberrant paracrine signalling for bone remodelling underlies the mutant histone-driven giant cell tumour of bone"

#### Content

Supplementary Methods

Supplementary Figures and Legends 1-7

Supplementary Tables and Legends 1-4

Captions for Supplementary Data (Excel Spreadsheet) 1-5

References for Supplementary material

### Supplementary Methods

**Digital droplet PCR (ddPCR) on plasma samples.** Blood samples were processed for extraction of cfDNA and DNA was analysed using ddPCR according to the protocol previously described (1). The presence of the canonical hTERT inactivating promoter mutation was detected using the BioRAD ddPCR EXD Assay TERT C228T_88, Has (dHsaEXD20945488, 12003908). The presence of the G34W mutation was detected using specific primers (G34W Fw 5’ AAGCAACTGGCTACAAAA 3’, G34W Rev 5’ TGGATACATACAAGAGAGA 3’) and probe (5’ CCTCTACTGGAGGGGTGAAGAAA 3’).

### Supplementary Figures and Legends

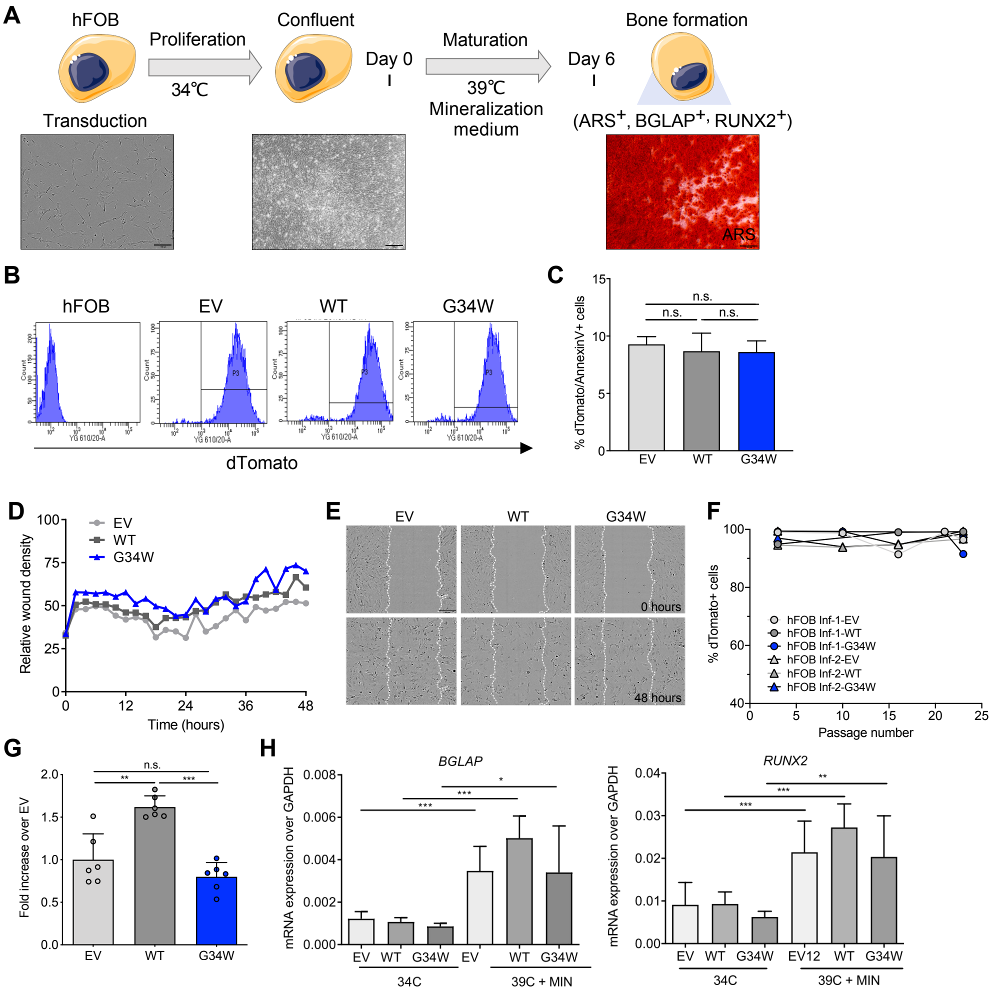


**Supplementary Figure 1. Stable expression of H3.3$^{G34W}$ in hFOB alters bone formation.**

**A.** Schema of timeline of hFOB differentiation to mature osteoblasts and representative bright field images of hFOB showing bone formation by alizarin red staining (ARS). **B.** Percentage of dTomato-positive hFOBs transduced with H3.3$^{WT}$, H3.3$^{G34W}$ and empty vector (EV) and assessed by flow cytometry after FACS sorting. **C.** Apoptosis of hFOB cultured at 34°C; 4 replicates. **D-E.** Wound healing assay: (D) quantification of the relative 'wound healing' assessed by Incucyte and (E) representative phase contrast images; 8 replicates, in 2 independent infections. **F.** Number of dTomato-positive hFOB in culture over time, in two independent infections (Inf-1 and Inf-2). All assays were performed using cells that were >90% dTomato-positive. **G.** Quantification of mineralisation of hFOB after 6 days of differentiation assessed by OsteoImage assay; 6 replicates. **H.** Gene expression of osteoblast genes *BGLAP* and *RUNX2* in hFOB at 34°C and after differentiation at 39°C in the presence of mineralisation medium (MIN) for 6 days; 2 independent experiments, 3 replicates per experiment.

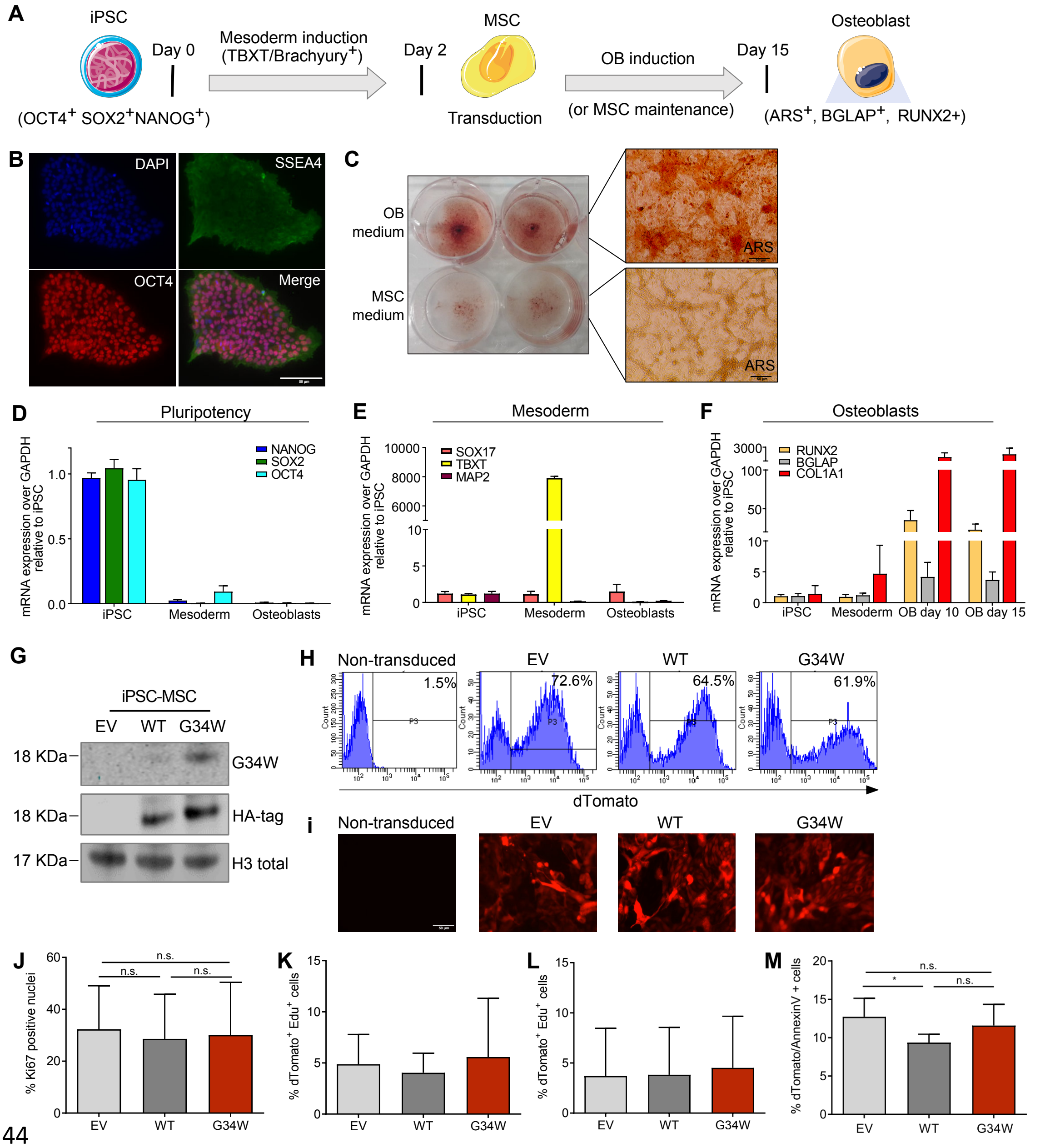


**Supplementary Figure 2. Stable expression of H3.3$^{G34W}$ in iPSC-derived MSCs.**

**A.** Schematic timeline of differentiation of iPSCs to mesenchymal stem cells (MSC) via mesoderm induction, and then either maintained as MSC in MSC maintenance medium (MSC medium) or induced to mature into mineralised osteoblasts in osteoblast-inducing medium (OB medium). iPSCs were differentiated in MSC medium for 48 hours, transduced with lentiviral particles containing dTomato empty vector (EV), H3.3$^{WT}$ or H3.3$^{G34W}$ and then differentiated to osteoblasts. **B.** Analysis of the pluripotency markers OCT4 and SSEA4 expression in iPSCs using immunofluorescence. **C.** Bright field photomicrographs of calcium deposition detection by Alizarin Red staining (ARS) in non-transduced iPSCs on day 15 of differentiation to osteoblasts in OB medium, and in cells maintained in MSC maintenance medium. **D-F.** (D) Expression of genes of pluripotency (*NANOG*, *SOX2*, *OCT4*) is high in iPSC cells and suppressed during mesoderm induction and OB differentiation. (E) Expression of mesoderm-specific genes (*TBXT*) is induced upon mesoderm induction, whereas expression of *SOX17* (marker of endoderm differentiation) and *MAP2* (marker of ectoderm differentiation) is suppressed. (F) Osteoblast-specific genes (*RUNX2*, *BGLAP*, *COL1A1*) are induced following osteoblast differentiation in a time-dependent manner; qPCR results, 3 replicates. **G.** Western blot: validation of $H3.3^{WT}$-HA and $H3.3^{G34W}$-HA overexpression on acid-extracted histone preparations of iPSC-derived MSCs. **H-I.** (H) Percentage of dTomato-positive iPSC-derived MSC as assessed by flow cytometry and (I) fluorescence microscopy 48 hours after transduction, 20X magnification. **J.** Number of Ki67-positive nuclei by immunofluoresence in iPSC-derived MSC differentiated osteoblasts (3 independent transductions, each with 2 replicates). **K-L.** EdU proliferation assay by flow cytometry: number of dTomato+EdU-positive cells on day 2 of (K) MSC maintenance in MSC maintenance medium and (L) osteoblast differentiation; 3 replicates. **M.** Apoptosis of iPSC-derived MSCs on day 4 of osteoblast differentiation assessed by AnnexinV-PI staining; 2 independent transductions, each with 3 replicates.

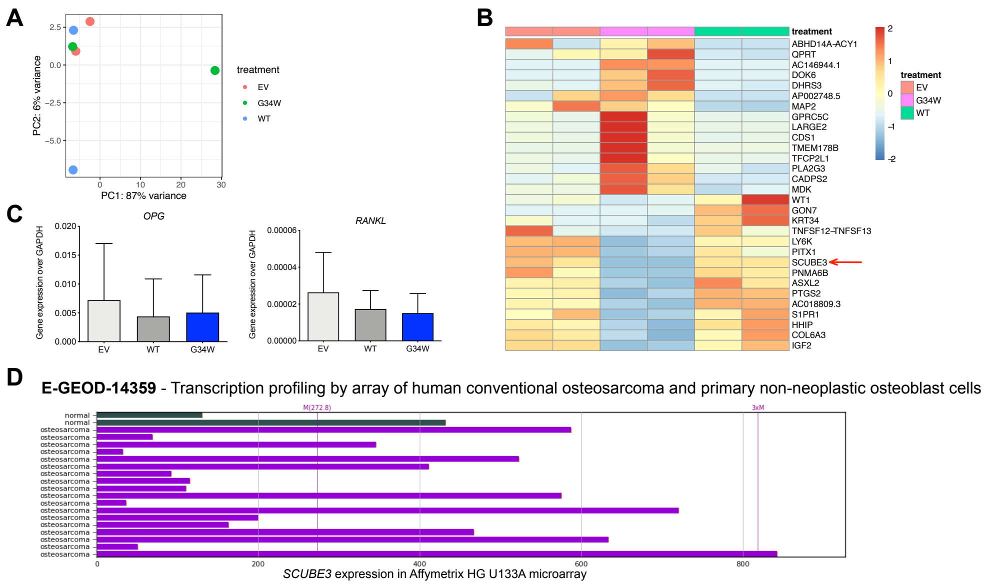


**Supplementary Figure 3. RNA-sequencing of hFOB, expression of *SCUBE3*.**

**A.** Principal component analysis (PCA) of RNA-seq data of hFOB performed on rlog transformed expression values. **B.** Heatmaps show statistically significant differences in gene expression, including *SCUBE3*, between wild type (WT) and G34W hFOB samples (selection of genes, Independent Hypothesis Weighting (IHW)-adjusted p-value < 0.001). A red arrow indicates *SCUBE3*. **C.** Expression of *OPG* and *RANKL* by qPCR in hFOB transfectants cultured at 34ºC. Data represent 6 experiments from 4 independent infections. **D.** Analysis of publicly available gene expression datasets reveal expression of *SCUBE3* in normal human osteoblast and human osteosarcoma samples (GSE14359(2)) as visualised on the BioGPS portal (3).

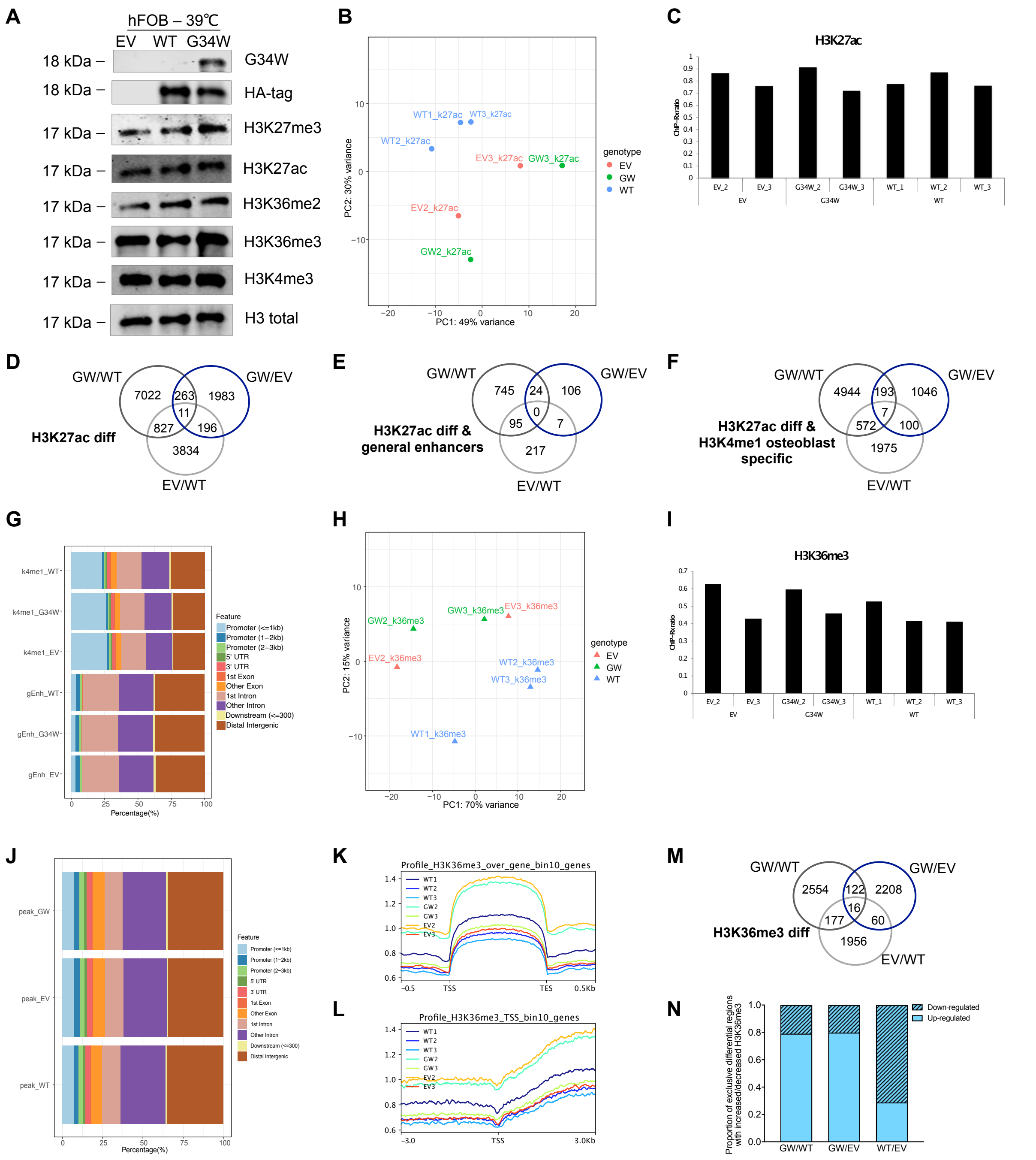


**Supplementary Figure 4. H3K27ac and H3K36me3 Chromatin Immuno Precipitation-sequencing normalised with an exogenous reference genome (ChIP-Rx) of hFOB.**

**A.** Validation of H3.3$^{WT}$-HA and H3.3$^{G34W}$-HA overexpression and total levels of histone marks (H3K27me3, H3K27ac, H3K36me2, H3K36me3, H3K4me3 and total H3) on acid-extracted histone preparations by western blot of hFOB after 3 days of differentiation at 39ºC. **B.** PCA of H3K27ac ChIP-seq data of hFOB. **C.** Ratio of sequencing reads mapping to human and drosophila genome in H3K27ac ChIP-seq, normalised by input. **D-F.** Venn diagram showing the number of differential H3K27ac peaks (D) overall, (E) intersected with general enhancers and (F) intersected with H3K4me1 osteoblast-specific genomic regions in hFOB. **G.** Distribution of genomic features of H3K27ac peaks intersected with H3K4me1 osteoblast-specific regions (top three bars) and intersected with general enhancers (bottom three bars) in hFOB. **H.** PCA of H3K36me3 ChIP-seq data of hFOB. **I.** Ratio of sequencing reads mapping to human and drosophila genome in H3K36me3 ChIP-seq, normalized by input. **J.** Distribution of genomic features of H3K36me3 peaks in hFOB. **K.** Positional profiles of H3K36me3 library around gene body. **L.** Positional profiles of H3K36me3 library around TSS. **M.** Venn diagram showing the number of differential H3K36me3 peaks in hFOB. **N.** Proportion of exclusive differential peaks showing increased (up-regulated) or decreased (down-regulated) H3K36me3 marks from pairwise comparison among H3.3$^{G34W}$ (GW), H3.3$^{WT}$ (WT) and EV.

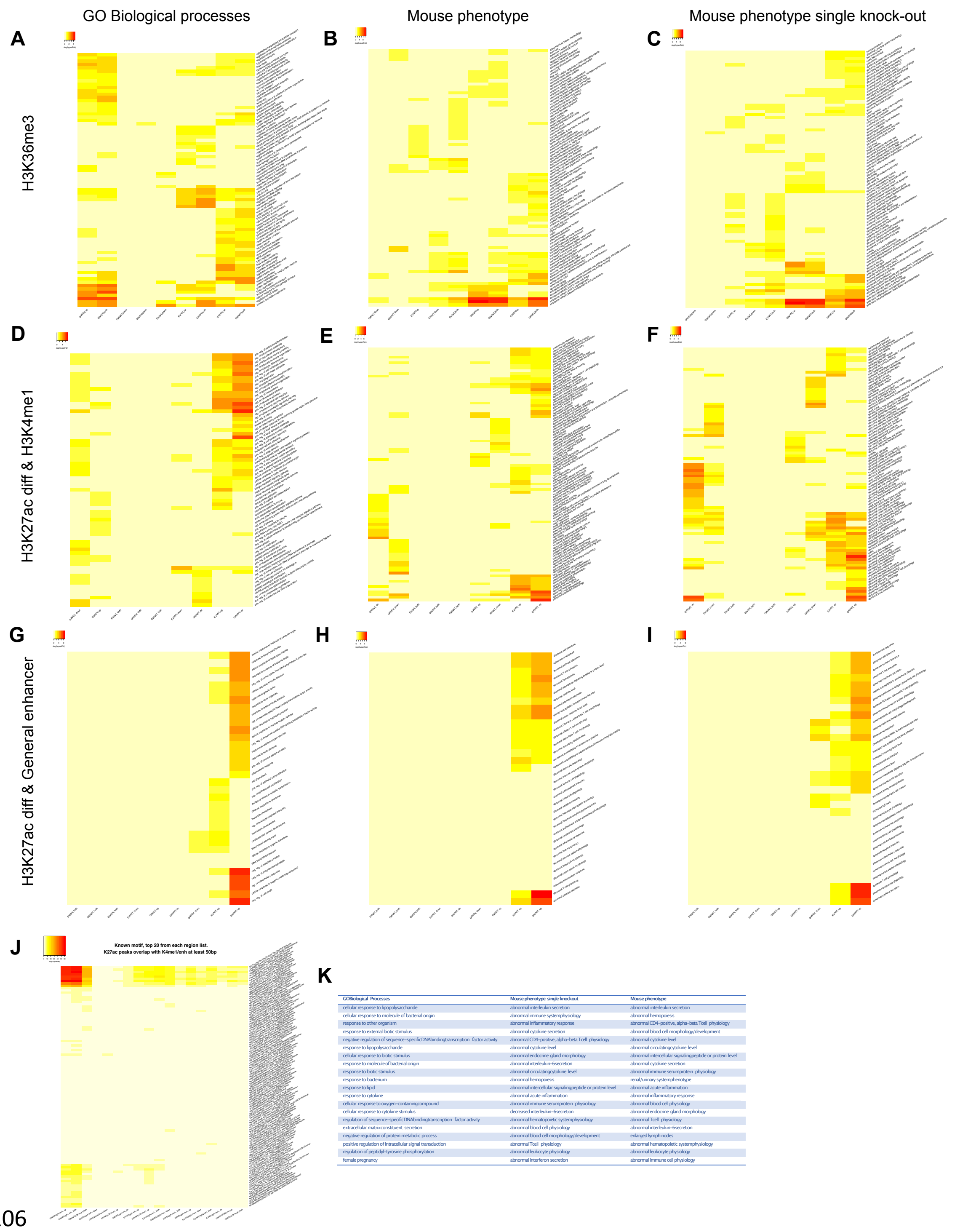


| GOBiological Processes | Mouse phenotype single knockout | Mouse phenotype |
|---|---|---|
| cellular response to lipopolysaccharide | abnormal interleukin secretion | abnormal interleukin secretion |
| cellular response to molecule of bacterial origin | abnormal immune systemphysiology | abnormal hemopoiesis |
| response to other organism | abnormal inflammatory response | abnormal CD4-positive, alpha-beta Tcell physiology |
| response to external biotic stimulus | abnormal cytokine secretion | abnormal blood cell morphology/development |
| negative regulation of sequence-specificDNAbindingtranscription factor activity | abnormal CD4-positive, alpha-beta Tcell physiology | abnormal cytokine level |
| response to lipopolysaccharide | abnormal cytokine level | abnormal circulatingcytokine level |
| cellular response to biotic stimulus | abnormal endocrine gland morphology | abnormal intercellular signalingpeptide or protein level |
| response to molecule of bacterial origin | abnormal interleukin-6secretion | abnormal cytokine secretion |
| response to biotic stimulus | abnormal circulatingcytokine level | abnormal immune serumprotein physiology |
| response to bacterium | abnormal hemopoiesis | renal/urinary systemphenotype |
| response to lipid | abnormal intercellular signalingpeptide or protein level | abnormal acute inflammation |
| response to cytokine | abnormal acute inflammation | abnormal inflammatory response |
| cellular response to oxygen-containingcompound | abnormal immune serumprotein physiology | abnormal blood cell physiology |
| cellular response to cytokine stimulus | decreased interleukin-6secretion | abnormal endocrine gland morphology |
| regulation of sequence-specificDNAbindingtranscription factor activity | abnormal hematopoietic systemphysiology | abnormal Tcell physiology |
| extracellular matrixconstituent secretion | abnormal blood cell physiology | abnormal interleukin-6secretion |
| negative regulation of protein metabolic process | abnormal blood cell morphology/development | enlarged lymph nodes |
| positive regulation of intracellular signal transduction | abnormal Tcell physiology | abnormal hematopoietic systemphysiology |
| regulation of peptidyl-tyrosine phosphorylation | abnormal leukocyte physiology | abnormal leukocyte physiology |
| female pregnancy | abnormal interferon secretion | abnormal immune cell physiology |

**Supplementary Figure 5. H3K27ac and H3K36me3 ChIP-Rx of hFOB: functional analysis. A-C.** Heatmaps of functional analysis for GO biological processes, mouse phenotype and mouse phenotype single knock-out for exclusive differential H3K36me3 peaks. **D-I.** Heatmaps of functional analysis for GO biological processes, mouse phenotype and mouse phenotype single knock-out for exclusive differential H3K27ac peaks intersected with (D-F) osteoblast-specific H3K4me1 genomic regions and (G-I) general enhancers in hFOB. **J.** Heatmap of TF binding motifs in exclusive differential H3K27ac peaks intersected with either osteoblast-specific H3K4me1 genomic regions or general enhancers in hFOB. **K.** Functional analysis of Ets and Ets-like motifs-containing differential H3K27ac peaks up-regulated exclusively in G34W vs WT intersected with general enhancers: Ets domains were sited close to genes involved in the immune response, cytokines and interleukin secretion.

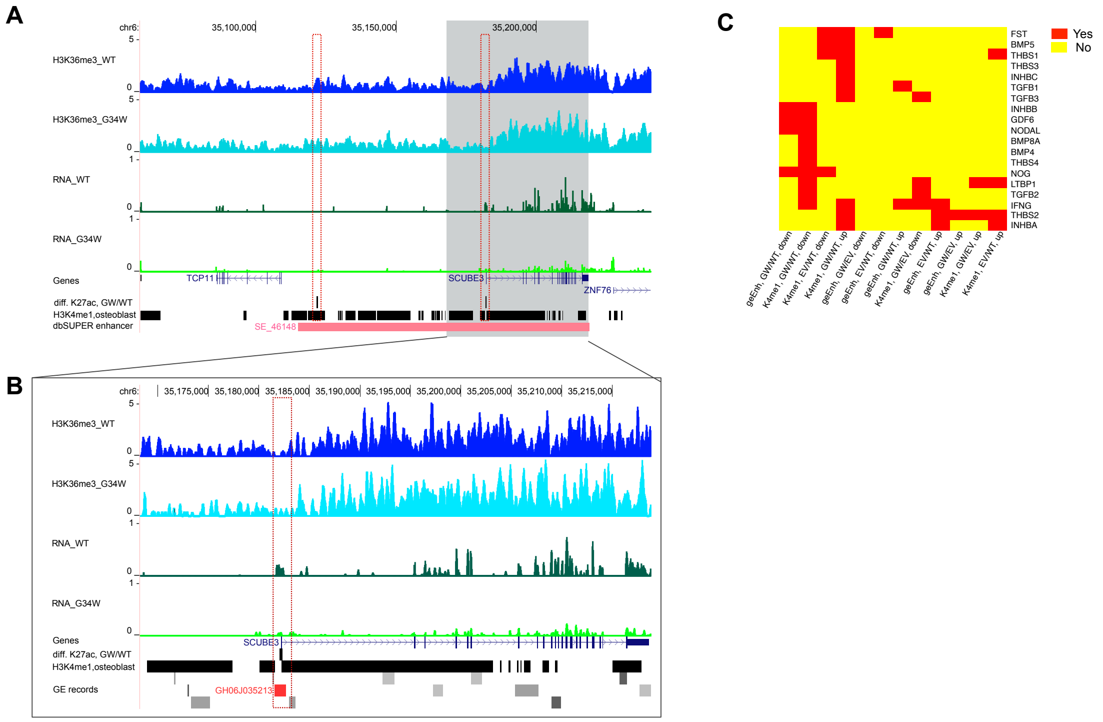


**Supplementary Figure 6. H3K36me3 alterations at the SCUBE3 locus and altered secreted genes.**

**A.** H3K36me3 modifications (top pair of tracks) at the *SCUBE* genomic locus and RNA expression (3rd and 4th tracks) in H3.3$^{WT}$ and H3.3$^{G34W}$ -hFOB. Differential H3K36me3 peaks from H3.3$^{G34W}$(GW) vs H3.3$^{WT}$ (WT) are shown in black. The super-enhancer SE_46148 (reported in the dbSUPER database) is shown in red. Tracks show one representative replicate. **B.** As (F) but with an expanded view of the *SCUBE3* gene. The enhancer GH06J035213 (reported in the GE database) is shown in red. **C.** Heatmap of secreted genes in the TGF−beta family with exclusive differential H3K27ac peaks in hFOB.

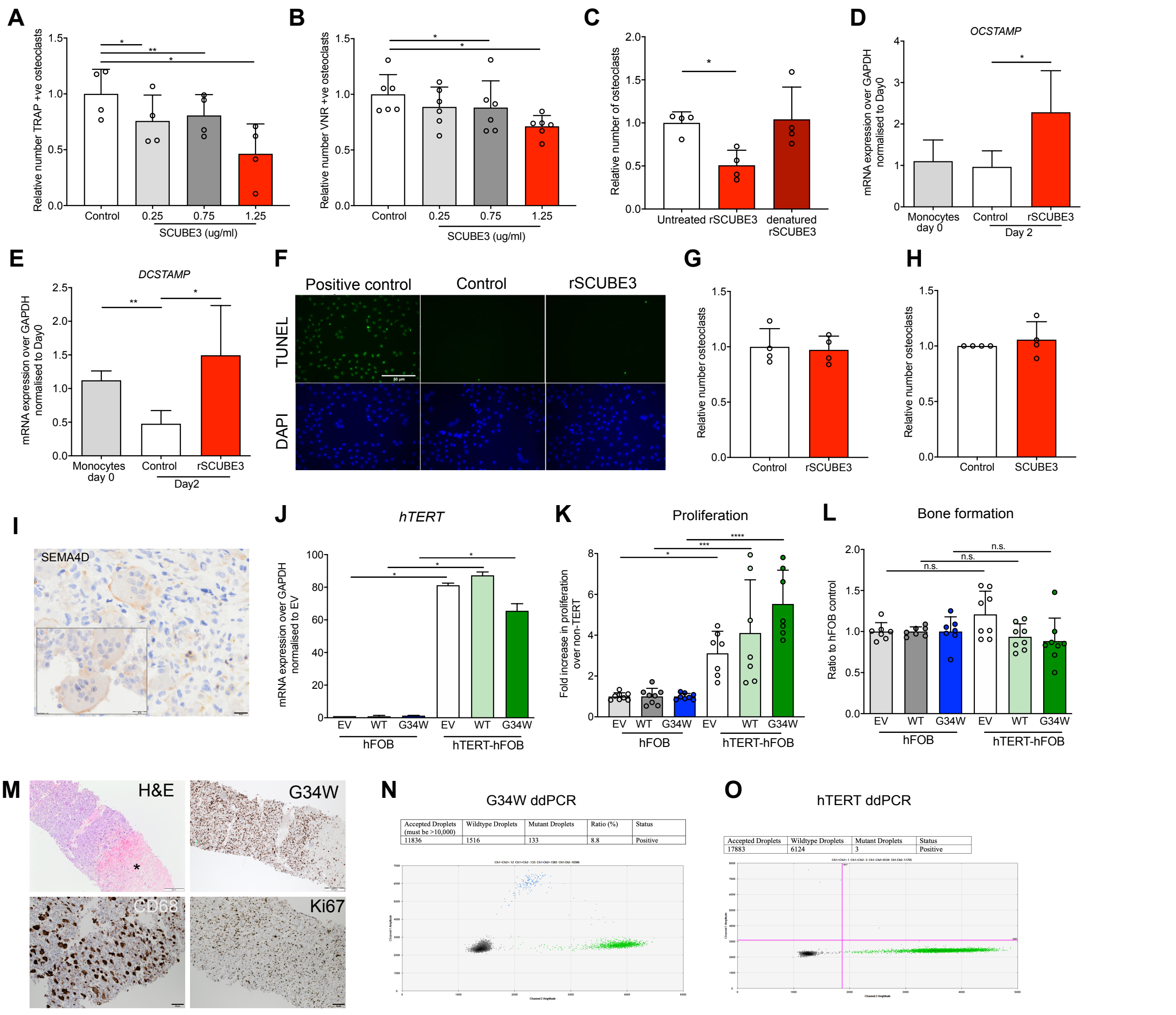


**Supplementary Figure 7. Role of SCUBE3 and SEMA4D in benign and malignant GCT.**

**A-B.** Number of (A) TRAP-positive and (B) VNR-positive osteoclasts on day 9 of differentiation in the presence of increasing concentrations of rSCUBE3; 4 and 6 osteoclasts preparations respectively. **C.** Treatment with 1.25 μg/ml rSCUBE3 denatured by heat inactivation does not reduce the number of osteoclasts formed; 4 osteoclasts preparations. **D-E.** rSCUBE3 alters the expression of genes involved in osteoclast fusion, *OCSTAMP* and *DCSTAMP*, assessed by qPCR, in untreated monocytes (day 0) and in osteoclasts differentiated from monocytes for 2 days in the absence or presence of 1.25 μg/ml rSCUBE3. Data represent 4 osteoclast preparations. **F.** Fluorescent images of TUNEL staining of osteoclasts at day 9 of differentiation, following differentiation in the presence of 1.25 μg/ml rSCUBE3; 3 osteoclasts preparations **G-H.** Survival of mature (day 8) osteoclasts treated for 48 hours with 1.25 μg/ml rSCUBE3 or vehicle: number of TRAP- (G) and VNR- (H) positive osteoclasts at day 10; 4 preparations. **I.** Photomicrograph of GCT showing immunoreactivity of SEMA4D depicting the membrane of osteoclasts; 40X magnification. **J.** qPCR analysis of expression of h*TERT* in hFOB transfectants before (hFOB) and after over-expression of hTERT (hTERT-hFOB). Data represent 2 replicates per condition. **K.** Proliferation of hFOB and hTERT-hFOB transfectants as assessed by Presto Blue assay after 7 days of proliferation at 34°C; data represent 7-8 replicates. **L.** Bone formation in hFOB and hTERT-hFOB transfectants on day 6 of differentiation as assessed by the Osteoimage assay; data represent 7-8 replicates. **M.** Photomicrograph of an osteoclast-rich G34W-mutant malignant GCT pre-treatment (histology of tumour shown in **Figure 5G**), showing necrosis (asterisk), presence of CD68-immunoreactive osteoclasts and atypical highly proliferative H3.3$^{G34W}$-mutant stromal cells (G34W, Ki67). **N-O.** Post-denosumab treatment (of tumour in **Figure 5G**) showing detection of G34W (N) and hTERT promoter (O) mutations in the circulating cell-free tumour DNA (cfDNA) using digital droplet PCR. This demonstrates that growth of the G34W-mutant tumour cells are independent of osteoclasts. Mutant G34W molecules in cfDNA have not been detected in benign GCT other than when associated with fracture (1).

### Supplementary Tables

**Supplementary Table 2.** Cell line authentication by STR (short tandem repeat) profiling of hFOB used in this study (report June 2020).

| | | UCL profiles | | Database profiles | |
|---|---|---|---|---|---|
| Cell line | Marker | Allele 1 | Allele 2 | Allele 1 | Allele 2 |
| **hFOB** | AMEL | X | X | X | X |
| | CSF1PO | 10 | 13 | 10 | 13 |
| | D13S317 | 11 | 12 | 11 | 12 |
| | D16S539 | 9 | 13 | 9 | 13 |
| | D18S51 | 10 | 17 | 10 | 17 |
| | D21S11 | 29 | 32.2 | 29 | 32.2 |
| | D3S1358 | 17 | 18 | 17 | 18 |
| | D5S818 | 11 | 12 | 11 | 12 |
| | D7S820 | 8 | 10 | 8 | 10 |
| | D8S1179 | 10 | 14 | 10 | 14 |
| | FGA | 19 | 22 | 19 | 22 |
| | Penta D | 9 | 13 | 9 | 13 |
| | Penta E | 8 | 11 | 8 | 11 |
| | TH01 | 7 | 9.3 | 7 | 9.3 |
| | TPOX | 11 | 11 | 11 | 11 |
| | vWA | 16 | 18 | 16 | 18 |

**Supplementary Table 3.** List of primers used for qPCR.

| | Gene | Primer sequences (5'-3') |
|---|---|---|
| **Control gene** | *GAPDH*_Fw | GAAGGTGAAGGTCGGAGTCA |
| | *GAPDH*_Rev | GAAGATGGTGATGGGATTTC |
| **Pluripotency markers** | *NANOG*_Fw | AACTGGCCGAAGAATAGCAA |
| | *NANOG*_Rev | TGCACCAGGTCTGAGTGTTC |
| | *OCT4*_F w | CCTCACTTCACTGCACTTGTA |
| | *OCT4*_Rev | CAGGTTTTCTTTCCCTAGCT |
| | *SOX2*_Fw | ATGTCCCAGCACTACCAGAG |
| | *SOX2*_Rev | GCACCCCTCCCATTTCCC |
| **Endoderm layer** | *SOX17*_FW | TGTTCAGAGATTTGTTTCCCATAG |
| | *SOX17*_RV | ACACACCCAGGACAACATTTC |
| **Ectoderm layer** | *MAP2*-Fw | CCACCTGAGATTAAGGATCA |
| | *MAP2*-Rev | GGCTTACTTTGCTTCTCTGA |
| **Mesoderm layer** | *TBXT*_Fw | CCCGTCTCCTTCAGCAAAGTC |
| | *TBXT*_Rev | TGGATTCGAGGCTCATACTTATGC |
| **Osteoblast differentiation** | *BGLAP*_Fw | AATCCGGACTGTGACGAGTT |
| | *BGLAP*_Rev | GGCAAGGGGAAGAGGAAAGA |
| | *RUNX2*_Fw | CTGTGGTTACTGTCATGGCG |
| | *RUNX2*_Rev | AGGTAGCTACTTGGGGAGGA |
| | *COL1A1*_Fw | GTGCTAAAGGTGCCAATGGT |
| | *COL1A1*_Rev | CTCCTCGCTTTCCTTCCTCT |
| | *SCUBE*_Fw | GTATGCTGGTTGTCGCTGAG |
| | *SCUBE*_Rev | GGTTGTGTGCATGACTGTGT |
| **hTERT** | *hTERT*_Fw | GCCGATTGTGAACATGGACTACG |
| | *hTERT*_Rev | GCTCGTAGTTGAGCACGCTGAA |
| **GCT human samples** | *SCUBE3*_70bp_Fw | GTGATGACACAGAGCAGGGT |
| | *SCUBE3*_70bp_Rev | CACAGGTCTCGATGCATGTCT |
| | *GAPDH*_78pb_Fw | CATACCAGGAAATGAGCTTGACAA |
| | *GAPDH*_78pb_Rev | ACACCCACTCCTCCACCTTTG |

**Supplementary Table 4.** List of antibodies used (western blot, IF, IHC, ChIP).

| Protein | Use | kDa | Brand | Cat. Number | Species | Clonality | Dilution |
|---|---|---|---|---|---|---|---|
| H3 Total | WB | 17 | Abcam | Ab 1791 | Rabbit | Polyclonal | 1:1000 |
| H3K27me3 | WB | 17 | Millipore | 07-449 | Rabbit | Polyclonal | 1:2000 |
| H3K36me3 | WB, ChIP | 17 | Abcam | Ab9050 | Rabbit | Polyclonal | 1:1000 |
| H3K36me2 | WB | 17 | CST | 9758 | Rabbit | Polyclonal | 1:1000 |
| H3K27ac | WB, ChIP | 17 | Abcam | Ab4729 | Rabbit | Polyclonal | 1:1000 |
| H3K4me3 | WB | 17 | Abcam | Ab8580 | Rabbit | Polyclonal | 1:1000 |
| *B-actin* | WB | 42 | Sigma | A5441 | Mouse | Monoclonal | 1:5000 |
| HA-tag | WB | | Abcam | Ab9110 | Rabbit | Polyclonal | 1:1000 |
| H3.3 G34W | WB, IHC | | RevMab Biosciences | 31-1145-00 | Rabbit | Clone RM263 | 1:250 |
| Anti-mouse Alexa Fluor 594 | IF | | Thermo Scientific | | Mouse | | 1:500 |
| Goat anti-Rabbit Alexa Fluor 488 | IF | | Thermo Scientific | A-11034 | Rabbit | Polyclonal | 1:500 |
| IRDye 800CW Goat anti-Rabbit | WB | | Thermo Scientific | SA5-35571 | Rabbit | Polyclonal | 1:5000 |
| IRDye 680CW Goat anti-Mouse | WB | | Thermo Scientific | SA5-35518 | Mouse | Polyclonal | 1:5000 |
| Ki67 | IF | | Abcam | Ab16667 | Rabbit | Clone SP6 | 1:200 |
| Ki67 | IHC | | Leica | PA0118 | Mouse | MM1 | Ready to use |
| CD68 | IHC | | Leica | PA0273 | Mouse | 514H12 | Ready to use |

*WB, Western blot.*

### Captions for Supplementary Data

**Supplementary Data 1 (Excel spreadsheet):**
**Supplementary_Data_1_hFOB_EV-WT_G34W_RNAseq_DEG.** List of DEGs with a p-adj value <0.05 identified by RNA-sequencing of hFOB (**Figure 2**): comparisons of G34W versus WT, G34W versus EV, EV versus WT.

**Supplementary Data 2 (Excel spreadsheet):**
**Supplementary_Data_2_H3K27ac_Pathway_Analysis_GWvsWT_all.** List of Gene Ontology (GOBP), mouse phenotype (MousePh) and mouse phenotype single knock out (MousePhSKO) analysis of differential H3K27ac peaks up- or down-regulated in G34W vs WT, intersected with osteoblast-specific H3K4me1 enhancers and general enhancers.

**Supplementary Data 3 (Excel spreadsheet):**
**Supplementary_Data_3_MatrixHmap_diffH3K27ac_spec_ALL.** –log10(HyperFdrQ) values for heatmaps of functional analysis for GO biological processes (GOBP), mouse phenotype (MousePh) and mouse phenotype single knock-out (MousePhSKO) for exclusive differential H3K27ac peaks intersected with H3K4me1 osteoblast-specific genomic regions and general enhancers in hFOB.

**Supplementary Data 4 (Excel spreadsheet):**
**Supplementary_Data_4_H3K36me3_Pathway_Analysis_GWvsWT_all.** List of Gene Ontology (GOBP), mouse phenotype (MousePh) and mouse phenotype single knock out (MousePhSKO) analysis of differential H3K36me3 peaks up- or down-regulated in G34W vs WT.

**Supplementary Data 5 (Excel spreadsheet):**
**Supplementary_Data_5_MatrixHmap_diffH3K36me3_spec_ALL.** –log10(HyperFdrQ) values for heatmaps of functional analysis for GO biological processes (GOBP), mouse phenotype (MousePh) and mouse phenotype single knock-out (MousePhSKO) for exclusive differential H3K36me3 peaks in hFOB.
